# HSP90-mediated stress resilience in male gametophyte of *Arabidopsis thaliana*

**DOI:** 10.1101/2025.06.21.660849

**Authors:** Zahra Kahrizi, Christos Michailidis, Karel Raabe, Jan Fíla, Božena Klodová, Jiří Rudolf, Petra Procházková Schrumpfová, David Honys, Sotirios Fragkostefanakis

## Abstract

Despite the accumulation of protective heat shock proteins (HSPs) during the male gametophyte development, pollen grains are highly sensitive to elevated temperatures. We performed transcriptomic analysis of five pollen developmental stages isolated from plants under normal and heat stress (H) conditions; uni-nuclear (UN), early bi-cellular (EB), bi-cellular (BC), tri-cellular (TC), and mature pollen (MPG). We show that the majority of genes that are up- or down-regulated under HS are specific for each stage, except BC stage that also exhibited the highest number of differentially expressed genes (DEGs) (>4000). This emphasize a complex, stage-dependent heat stress response, possibly dependent on HSP levels. Additionally, promoter motif analysis revealed that heat shock elements (HSEs) exhibit a stage-specific pattern of enrichment, which peaked in MP. To explore stage-specific influences of HSP90s in pollen development, we characterized a knockdown RNAi line, under normal and stress conditions in early- and late-stage RNAi lines using stage-specific promoters pJASON (JA90R) and pLAT52 (L90R). The *hsp90* background leads to lower germination rate in both RNAi lines that is more pronounced under heat stress caused by significant alterations in heat stress control via impaired ABA signalling or ER stress response. Heat stress conditions also lead to a high percentage of nuclei shape and orientation defects in the L90R line pollen, pointing to the higher sensitivity of late stage development. We show this defect is linked to the down-regulation of DNA metabolism genes. Our complex dataset provides insight into stage-specific stress response on the level of single cell undergoing developmental changes.

## Introduction

High temperature events result in reduction in plant fertilization success (Folsom et al. 2014; Zenda et al. 2021; Begcy et al. 2019; Nadakuduti et al. 2023), where male gametophyte development is one of the most heat-sensitive developmental phase in plant reproduction (De Storme and Geelen 2020; Begcy et al. 2019; Chaturvedi et al. 2021).

In *Arabidopsis thaliana*, male gametophyte development consists of two phases: microsporogenesis and microgametogenesis. During the first phase, a pollen mother cell (microsporocyte) undergoes meiosis, producing four microspores. In the second phase, each microspore undergoes two rounds of mitotic divisions, pollen mitosis I (PMI) and II (PMII), to form tri-cellular structure the mature pollen grains (MPGs) (Oh et al., 2020; Hafidh and Honys 2021; Li et al., 2025). The PMI division is asymmetric, yielding a small generative cell that is later engulfed into the cytoplasm of much larger vegetative cell. In PMII, the generative cell divides to two sperm cells. Hence, the stages of pollen development reflect the number of the microspore nuclei: uninuclear (UN), early bi-cellular (EB), bi-cellular (BC) and tri-cellular (TC) pollen and finally the dehydrated mature pollen grain (MPG). After pollination, the MPG rehydrates and germinates into pollen tube, a transport vehicle that deliver sperm cells through the pistil to fertilize the female embryo sac (McCormick 2004).

In response to higher temperatures, plants activate heat shock transcription factors (HSFs) to regulate the expression of Heat shock protein (HSP) genes (Ohama et al., 2017; Bakary et al., 2024; Fragkostefanakis et al., 2025). HSPs are then engaged in various biological processes during environmental or endogenous stresses by protecting protein homeostasis and cellular integrity, obstructing cell death signalling, and facilitating DNA repair (Sottile and Nadin, 2018; Dubrez et al., 2020).

HSPs, many of which act as molecular chaperones, are categorized into several groups based on their molecular weight, namely HSP100, HSP90, HSP70, HSP60, and the small HSP family (sHSP), each with unique functions in the plant life cycle. The group of HSP90s is characterized as proteins acting as dynamic molecular scaffolds that ensure stability and function of various signalling pathways that regulate plant growth and response to environmental cues (Zhou et al., 2020).

Seven HSP90 genes have been have been identified within the genome of *A. thaliana*: *AtHSP90-1* (AT5G52640), *AtHSP90-2* (AT5G56030), *AtHSP90-3* (AT5G56010), and *AtHSP90-4* (AT5G56000) proteins that are localized in the cytoplasm, whereas *AtHSP90-5* (AT2G04030), *AtHSP90-6* (AT3G07770), and *AtHSP90-7* (AT4G24190) are localized in chloroplasts, mitochondria, and endoplasmic reticulum (ER), respectively (Milioni, et al., 1997). In general, some *HSP90* genes are constitutively expressed, while others are induced by stress stimuli. This regulation of *HSP90s* expression is fine-tuned and its perturbations affect growth and development as well as general stress resilience via HSP90s impacting mechanisms of cell division, development, and flowering time (Sangster et al., 2008). For example, *HSP90.1*, *HSP90.2*, *HSP90.3*, and *HSP90.4* control flowering time by interacting with transcription factors that regulate flowering on the epigenetic level in a temperature-dependent manner, contributing to the coordination of development based on environmental signals (Margaritopoulou, et al., 2016; Zhou et al., 2020). Additionally, HSP90-1 and HSP90-2 control auxin-regulated flowering time by interacting with auxin response factor, (APRF1) (Isaioglou et al., 2024).

In another study, overexpression of *HSP90.2*, *HSP90.5*, and *HSP90.7* resulted in decreased activity of antioxidant enzymes, indicating increased protection and enhanced drought tolerance in *A. thaliana* plants (Song et al., 2009). Interestingly, knockout of *HSP90.1* or overexpression of *HSP90.3* both led to reduced thermotolerance in *A. thaliana* (Xu et al., 2012; Song et al., 2012). This contrasting effect highlights the functional diversity of HSP90 genes in stress responses, which is supported by the dual role of HSP90 proteins as essential molecular chaperones for thermotolerance and mediators of HSF activity (Prodromou, 2016).

At optimal temperatures, constitutively expressed HSFs remain inactive due to their interaction with HSP70 and HSP90 chaperones (Hahn et al., 2011). When temperature increases, the accumulation of unfolded proteins together with changes in redox state, induction of Ca^2+^ signalling and changes in post-translational modification, lead to the release of HSFs from HSPs, which have a higher affinity for unfolded proteins (Mesihovic et al., 2022; Bakary et al., 2024). Afterwards, HSP90s are among the hundreds of genes transcriptionally induced by HSFs, so their production improves overall protein folding mechanisms, cellular survival and recovery from heat stress. In tomato, HSP90 and HSP70 proteins interact with key HSFs like HsfA directly, influencing its DNA-binding activity and stability (Hanh et all 2011; Tiwari et al., 2020). In summary, HSP90 plays positive and negative roles in modulating both abundance and activity of HSFs, depending on the cellular context (Hahn et al., 2011). Furthermore, inhibitors of HSP90 activity (geldanamycin) lead to increased HsfA2 expression in *A. thaliana* (Nishizawa-Yokoi et al., 2010). This suggests that *Arabidopsis* HSP90 also suppresses Hsf activity under non-stress conditions, acting as a regulatory checkpoint for heat-induced transcriptional responses.

Several HSP90s are induced during the early stages of pollen development in tomato (Fragkostefanakis et al. 2016). The accumulation of HSPs is considered a pro-active priming mechanism that pollen activates to compensate for the high sensitivity in case of an upcoming heat stress incident (Xie et al., 2022). Interestingly, while the HSF-dependent heat stress response is activated in pollen mother cells and microspores, the ER-related UPR which is dependent on ER-associated bZIP transcription factors is activated in the late stages of pollen development and has been shown to be essential for pollen germination and pollen tube growth (Fragkostefanakis et al., 2016). HSP90 is essential for pollen’s complex network of responses to environmental stressors, guaranteeing plant reproduction.

Currently the role of HSP90 in pollen development and heat stress response remains elusive. Here we present a complex set of data considering transcriptomic response of *A. thaliana* accession Col-0 male gametophyte to heat stress, described in Figure 1. To investigate the function of HSP90 across male gametophyte developmental stages and describe its importance for thermotolerance, we used RNA interference (RNAi) lines of selected HSP90 genes in *A. thaliana* that suppressed HSP90 expression either at early or late developmental stages. *A. thaliana* RNAi lines targeting both *HSP90-1* and *HSP90-3* simultaneously at distinct developmental stages were created. The *JASON* promoter, active during meiosis of pollen mother cell, was used to induce RNAi line in early-stage male gametophyte development (JA90R) (UN&EB). The *LAT52* promoter which directs gene expression in late-stage pollen development (MP) was used to generate RNAi line in late stages of male gametophyte (L90R). Under control and heat stress conditions, the L90R lines exhibited lower germination rate and significant percentage of impaired male germ unit structure compared to the wild-type. Under heat stress, unique and shared genes were impacted in each RNAi line, demonstrating the stage-specific effects of HSP90 deficiency under stress conditions. The HSP90 mutation causes an adverse context, resulting in changes in heat stress response during the late stages of male gametophyte development. It is especially pronounced under heat stress treatment in the RNAi line, where dysregulated genes involved in DNA repair, chromatin remodelling, and ER stress response lead to failure in an adaptive response of the male gametophyte that maintains cellular integrity. Taken together, we provide a unique insight into heat stress response on single-cell level, using a highly reduced organism that already adapted HSP accumulation as part of its life cycle.

**Figure 1.**
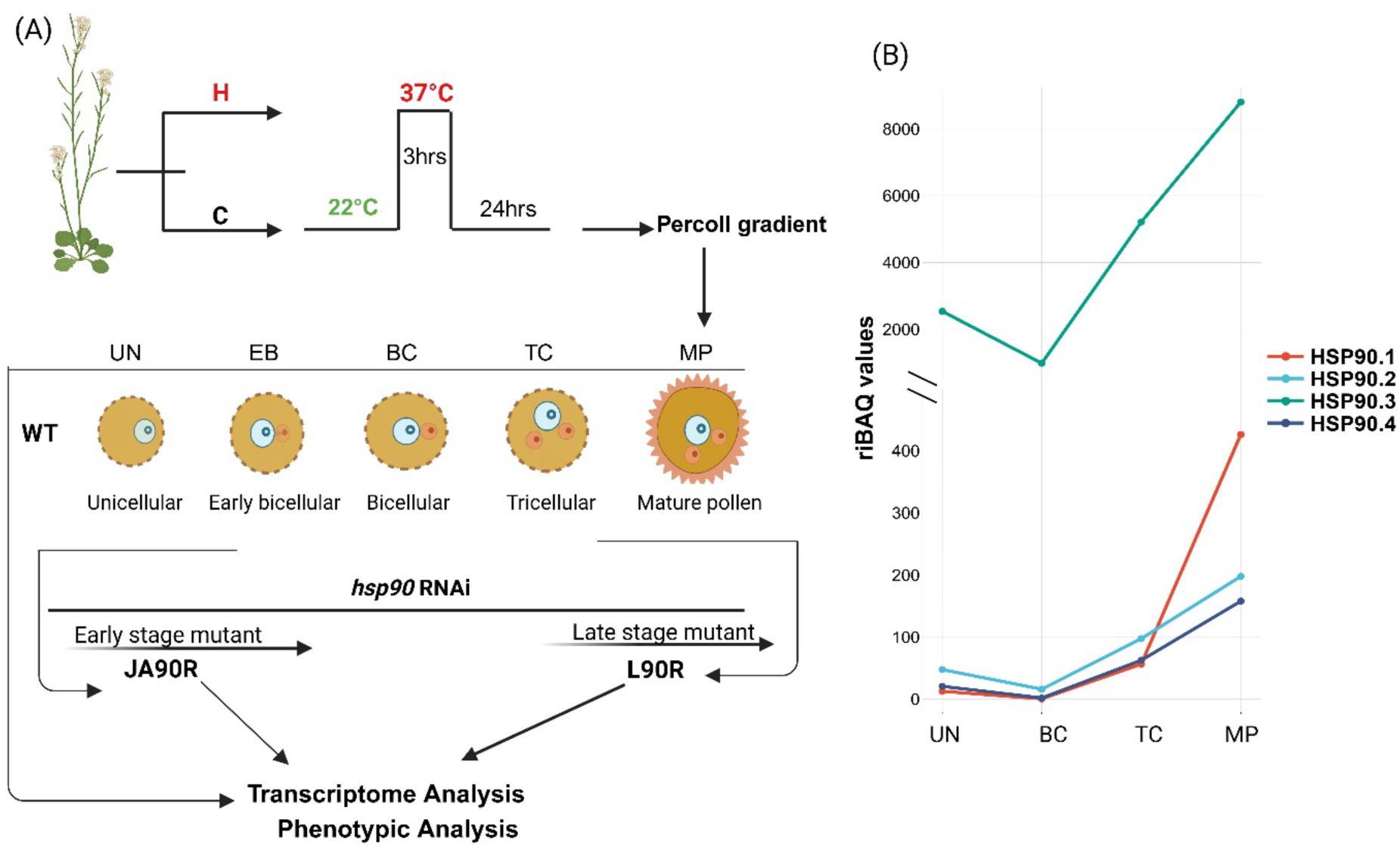
Representative image (**A**) showing the experimental setup, where pollen stages were isolated from *A. thaliana* Col-0 (WT) under control (C, 22 °C) and heat stress (H, 37 °C) conditions, followed by a 24- hour recovery period. Male gametophyte developmental stages included: Uni-cellular (UN), early bi- cellular (EB), bi-cellular (BC), tri-cellular (TC), mature pollen (MP) were isolated using a Percoll gradient for transcriptome analysis. RNAi lines targeting *HSP90-1* and *HSP90-3* were induced at early (JA90R) and late (L90R) stages using stage-specific promoters (regulated by *JASON* and *Lat52*). (**B**) The riBAQ values, based on our published proteome data (Klodová et al., 2023), highlight dynamic HSP90 expression, particularly with increased levels of HSP90-1 and HSP90-3 during late stages of pollen development under control conditions.

## Results

### Heat-stressed pollen shows stage-specific transcriptome changes

Pollen development has been best described in the model plant *A. thaliana* and stage-specific transcriptome changes accompanying the dynamic cellular, structural and metabolic changes have been reported (Wei & Ma, 2023). However, how elevated temperatures affect gene expression in a stage-specific manner is currently addressed in *A. thaliana.* Here, *A. thaliana* Col-0 plants at inflorescence stage were exposed to 37°C for three hours, while control plants were kept at 22 °C control condition. Pollen was isolated from the stressed and control plants, and fractionated into five distinct developmental stages, namely UN, EB, BC, TC and MP (Figure 2a). Transcriptome analysis by RNA-Seq revealed common and unique patterns of gene expression during the response to heat stress at individual stages. We did not observe a major difference between the number of up-regulated and down-regulated genes in each stage, however, heat-stressed pollen at BC stage exhibited the highest number of differentially expressed genes (DEGs) (> 4000 up-regulated genes) compared to the other stages where the numbers were in hundreds of DEGs (271-879 genes) (Figure 2b). For all stages except BC, the majority of genes that were up- or down-regulated appear to be specific for each stage (Figure 2c). Interestingly, we also identified genes that are induced by heat in one stage, but reduced in another stage and vice versa (Figure S1), suggesting a complex and largely stage-dependent regulation of heat stress response throughout *A. thaliana* male gametophyte development.

**Figure 2.**
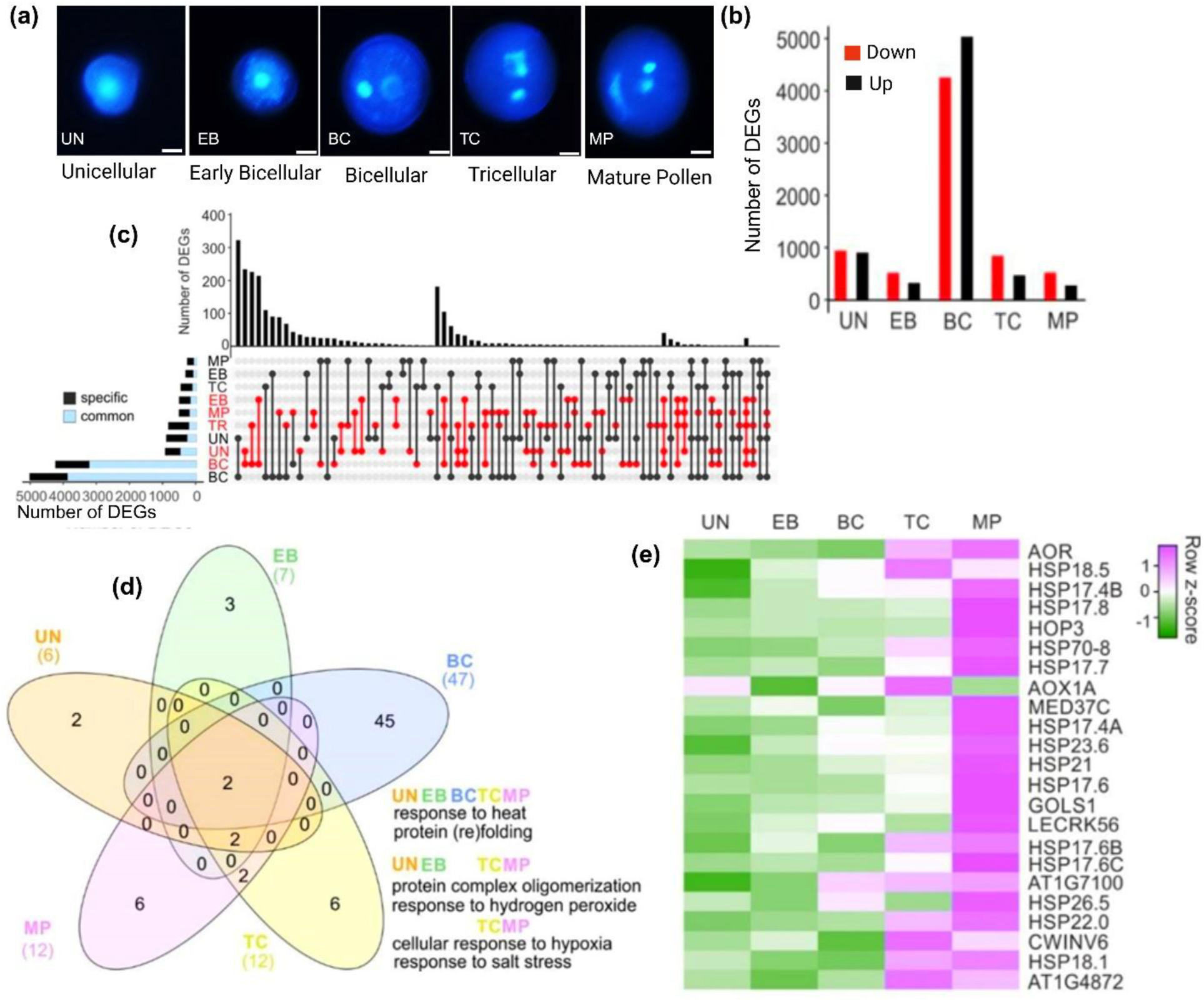
Pollen stage-specific transcriptome changes in response to heat stress in WT. (**a**) Representative images and illustrations of the isolated control pollen stages. (**b**) Number of differentially expressed genes (DEGs) (up-regulated, down-regulated in uni-cellular (UN): 890, 851, early bi-cellular (EB): 466, 271, bi-cellular (BC): 4206, 4980, tri-cellular (TC): 793, 419, and mature pollen (MP): 471, 691. Absolute Fold change > 1, p < 0.01). (**c**) UpSet plot showing the intersection of up- and down-regulated gene sets in different pollen stages. The number of genes that are up- or down-regulated specifically in one stage are indicated in the horizontal bars as fraction of the total. (**d**) Venn diagram showing the overlap of the functional gene ontology categories for heat stress induced genes in each pollen stage. Categories appearing in more than 2 stages are shown on the bottom right site of the panel. (**e**) Heat map showing the expression of a set containing 23 genes that are heat stress-induced in all pollen stages.

A gene ontology analysis was performed to examine whether this stage-specific gene regulation is related to specific biological functions. Two functional categories, namely “response to heat” and “protein (re) folding” were enriched in all stages (Figure 2d). In addition, “protein complex oligomerization” and “response to hydrogen peroxide” were enriched in all stages except the BC stage, while “cellular response to hypoxia” and “response to salt stress” were enriched only in TC and MP stages. In the BC stage, heat stress response involves a significant transcriptional regulation across diverse pathways, including RNA processing (snRNA, tRNA, rRNA) and surveillance systems, mitochondrial and chloroplast transcription, and protein modification processes (Supplementary data S1). Additionally, genes linked to meiosis, pollen development, and metabolic processes, such as carbohydrate, amine, and sulphur metabolism, are differentially expressed. In the MP stage, many up-regulated genes were related to endoplasmic reticulum (ER) stress response such as “ERAD pathway”, “ER unfolded protein response”, and “ER to Golgi vesicle-mediated transport” (Figure 2d). Among the >4000 heat stress induced genes, only 23 are up-regulated in all stages (Figure 2e). Among them, 13 belong to the highly induced small HSPs. Interestingly, the induction is stronger in later stages of pollen compared to early stages that all these genes show a much weaker induction, suggesting a difference in the amplitude of stress response among early and late stages for the core heat stress responsive genes.

The differences in the induction of sHSPs among the pollen stages in response to heat stress point to differences in proteostasis levels and possibly to different cellular requirements for HSPs. We checked the chaperone profile of cytosolic HSP90-, HSP70- and HSP20-coding genes in the different pollen stages under control and heat stress conditions, based on their cumulative transcript levels (Figure 3). We observed that the total levels of HSPs transcripts only weakly change in response to heat stress. However, the total levels peak in the UN stage for HSP90 and HSP70 genes, with a second accumulation at TC for HSP70 and MP for HSP90, respectively (Figure 3a-b). Among the four HSP90 members, HSP90-1 accumulates significantly in heat-stressed UN pollen, whereas in TC pollen, both HSP90-1 and HSP90-3 are expressed at substantially higher levels than the other two members under both conditions (Figure 3a). Our previously published proteome data shows the increased expression of HSP90-1 and HSP90-3 in the late stage of pollen development and lower expression in BC stage under control condition (Klodová et al., 2023) (Figure 1b). In contrast, HSP20s exhibit a strong accumulation in the EB stage under control conditions and show a strong accumulation in UN, BC and MP exposed to heat stress (Figure 3c).

**Figure 3.**
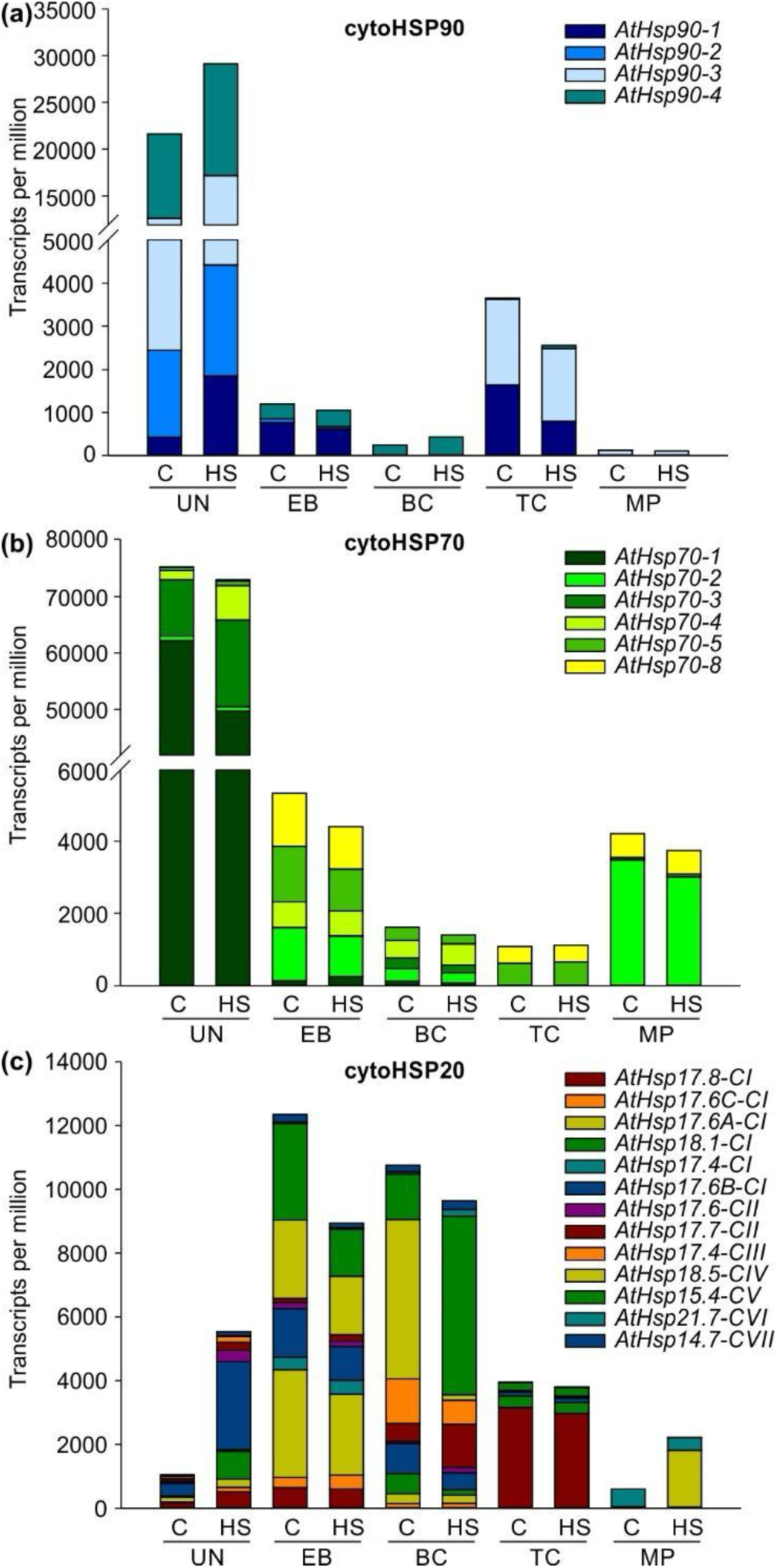
Cumulative expression of three families of HSPs during pollen development and heat stress response. Transcript levels of the indicated HSPs in five pollen stages under control (C) or heat stress (H) conditions. Only the genes coding for cytosolic/nuclear (**a**) HSP90, (**b**) HSP70, and (**c**) HSP20 are shown. UN: uni-cellular ; EB: early bi-cellular ; BC: bi-cellular ; TC: tri-cellular ; MP: mature pollen.

To test whether HSP transcript patterns reflect underlying cis-regulatory changes, we investigated the enrichment of various gene regulatory elements within the promoter regions from -1000 bp to +1000 bp relative to the translation start site (ATG) in the top-expressed genes at each pollen developmental stage using GOLEM program (Nevosád et al., 2025) (Figure S2). Our analysis revealed a slight increase in the number of genes containing heat shock elements (HSEs) in the promoters MP under heat stress compared to MP under control conditions. Specifically, the proportion of genes with an HSE motif located approximately -200 bp upstream of the ATG was higher in heat-stressed MP than in the control. In contrast, the frequency of HSE-containing genes in earlier stages - UN, EB, BC, and TC - showed minimal or no differences between control and heat stress conditions. The HSE enrichment profile diverged significantly from that of other responsive cis-elements associated with pollen development or stresses. For instance, the *telo*-box motif showed a clear increase in frequency under heat stress in BC but was not enriched in MP. Furthermore, no significant differential enrichment between control and heat stress conditions was observed for the DRE/CRT motif, TATA-box elements, or the *LAT52* promoter motif across male gametophyte stages.

In summary, we showed that the transcriptional response to heat is most pronounced in the BC stage, where, at the same time, the abundance of *HSP90* and *HSP70* transcripts is the lowest. This suggests that HSPs presence in the other stages could buffer the impact of stress and need to change the transcriptional profile. Furthermore, promoter motif analysis indicates that mature pollen possesses the responsive cis-regulatory architecture through HSEs, which likely facilitates its robust induction of small HSPs and other chaperones.

### HSP90 knockdown leads to defects in pollen phenotype and pollen germination

Based on our previous observations regarding the correlation of the HSP90 levels with the robustness of heat stress response, we wanted to see the fitness of pollen development under heat stress in low-HSP90 conditions. To specifically study the role of highly abundant HSP90s throughout the male gametophyte development, we utilised a 602 bp sequence of *HSP90-3* that has high similarity with *HSP90-1* sequence and used this sequence as RNAi target to down-regulate the two most abundant HSP90 members along the developmental stages. We created two RNAi lines that differ in the stage-specific promoter; JA90R active in early stages of male gametophyte development (under *JASON* promoter) and L90R active in late stages of pollen development (under *LAT52* promoter). The sequence was PCR-amplified and subsequently cloned into the GoldenBraid system in both sense and antisense orientations to form a hairpin RNA. This approach enabled to answer whether HSP90s’ role in early pollen development differs from its role in late stage of pollen development (Wielopolska et al, 2005; Bate & Twell, 1998; Storme & Geelen, 2011).

To confirm the knockdown RNAi lines, we checked the expression levels of *HSP90* genes in MP transcriptome of WT, L90R, and JA90R lines under control (C; 22 °C) and heat stress (H; 37 °C) conditions. Under heat stress, WT shows increased expression of several *HSP90* genes, whereas in L90R and JA90R lines, this transcriptomic response was significantly impaired (Figure 4a). The expression of specific *HSP90* genes in WT vs. RNAi lines shows decrease in *HSP90-1* and *HSP90-3* under control and heat stress conditions. This demonstrates that RNAi disrupts the target HSP90 expression levels and impairs their transcriptional response to heat stress (Figure 4b).

**Figure 4.**
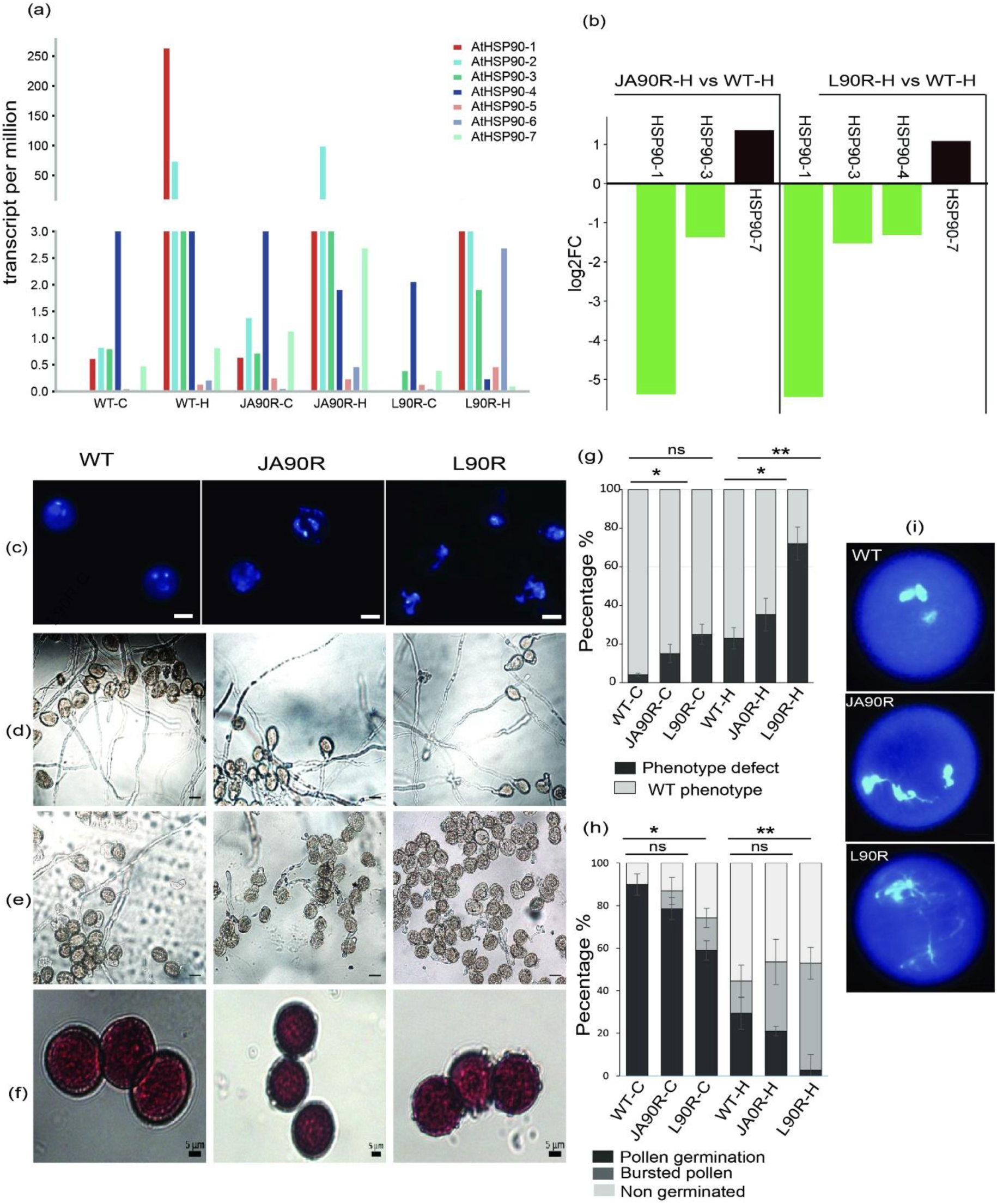
(**a**) Transcript levels of *HSP90* genes in mature pollen under control conditions (22°C; WT-C), and RNAi lines (L90R-C, JA90R-C) and under heat stress conditions (37 °C; WT-H), and RNAi lines (L90R-H, JA90R-H). (**b**) log2 fold change in HSP90 in WT vs L90R and JA90R under control (C) and heat stress conditions (H) in mature pollen transcriptome. (**c**) Pollen phenotype defects in WT, L90R and JA90R. The pollen phenotype defects in mature pollen in L90R and JA90R lines, mainly under heat stress conditions. Scale bar: 10 µm. (**d**) Pollen germination ratio in WT, L90R and JA90R under control (C) condition (22 °C) (e) Pollen germination ratio in WT, L90R and JA90R under heat stress (H) condition (37 °C) (f) Pollen viability test by alexander staining in WT, L90R and JA90R under heat stress (H) condition (37 °C) (**g**) Bar chart showing the percentage of pollen phenotype defects in WT, L90R, and JA90R lines at 22°C and 37°C. Gray bars represent WT phenotype pollen, Dark bars represent pollen with phenotype defects. Statistical significance is marked with * (p < 0.05) and ** (p < 0.01) by using Student’s t-tests. (**h**) Bar chart showing the percentage of pollen germination, burst pollen and non germinated pollen in WT, L90R, and JA90R lines at 22°C and 37°C. (**i**) The close up pollen phenotype defects in mature pollen in L90R and JA90R lines. Statistical significance is marked with * (p < 0.05) and ** (p < 0.01) by using Student’s t-tests.

Next, we hypothesized that decreased levels of HSP90 will lead to defects in pollen development and fitness. We screened MP from open flowers and observed defects in the L90R and JA90R lines under heat stress (Figure 4c-h). When compared to WT pollen, the L90R and JA90R lines displayed significant defect of impaired male germ unit (MGU) organisation phenotype, especially under heat stress. This argues for HSP90 knockdown affecting pollen viability and integrity more severely in the RNAi lines under stress conditions (Figure 4c, g). The quantified percentages of defective pollen in each line (WT, L90R, and JA90R) showed that at the control temperature (22°C), WT and JA90R exhibit minor increase in pollen defects, and L90R shows a slight rise in impaired MGU. However, under heat stress, L90R and JA90R RNAi lines show a significant increase in pollen MGU defects (64.8 % and 35.2 % respectively) compared to WT (23.1%) (Figure 4c, g). To further test RNAi line pollen viability, we performed a pollen thermotolerance assay by exposing buds to 37°C for three hours. After 24 hours, pollen viability and germination were tested. Alexander staining confirmed that pollen remained viable in WT, JA90R, and L90R under control conditions (Figure 4f). However, under heat stress, pollen germination defects increased especially in L90R, with 50.36% of pollen bursting, compared to 29.6% in WT and 20.96% in JA90R (Figure 4d, h). Taken together, we show here that disruption of HSP90 members throughout pollen development has the expected effect on pollen development and viability with the surprising effect of decreased HSP90 levels on the nucleus shape and MGU orientation after HS treatment.

### Down-regulation of DNA metabolism genes contributes to defects in MGU

To understand the observed phenotypic defects on molecular level, we analysed the transcriptomic distinct separation between control and heat-stressed samples, as well as between WT and RNAi lines (Figure 5a). The overlap of DEGs across the RNAi line-control condition comparisons showed 486 DEGs uniquely affected in L90R-C. Additionally, 214 genes were differentially expressed in JA90R-C and 24 genes differentially expressed across all comparisons (Figure 5b). Interestingly, L90R background has a set of core regulatory genes consistently expressed differentially in control and stress conditions (Figure 5c). Additionally, in L90R lines, next to down-regulation of *HSP90*, there was a marked downregulation of *HSP17, HSP18*, and *HSP70-5*, while *HSP70-3* was up-regulated.

**Figure 5.**
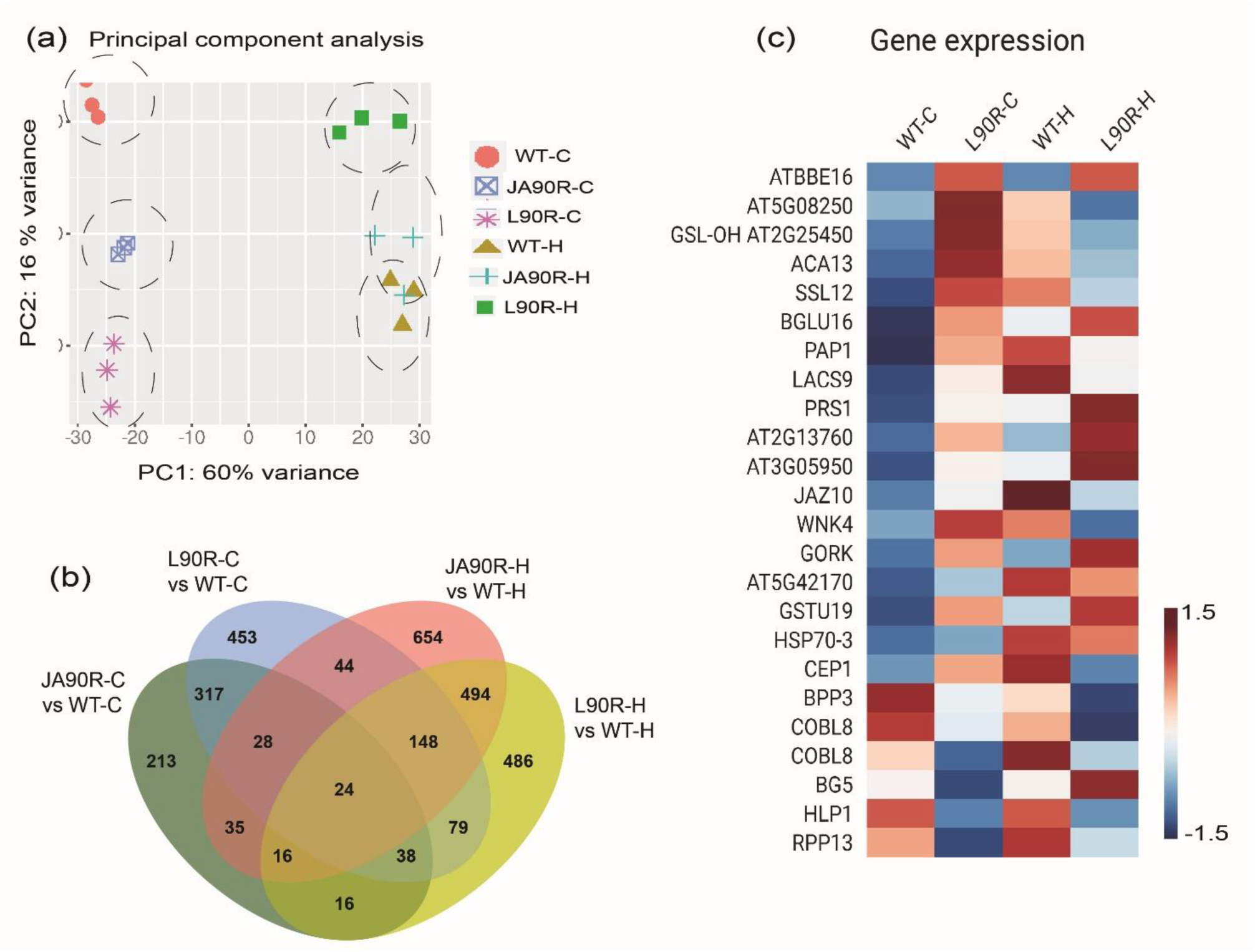
Differential gene expression analysis in pollen samples of WT, L90R, and JA90R lines (**a**) Principal Component Analysis (PCA) plot visualizing the variance in gene expression profiles between WT vs L90R, and JA90R lines under control (C) and heat stress conditions (H). PCA components account for 60% (PC1) or 16% (PC2) of the total variance. (**b**) Venn diagram illustrating the overlapping and unique differentially expressed genes (DEGs) between HSP90 knockdown RNAi lines (L90R and JA90R) and wild type (WT) under control (C) and heat stress (H) conditions. The numbers represent unique and shared DEGs across the four comparisons: L90R-C vs. WT-C, L90R-H vs. WT-H, JA90R-C vs. WT-C, and JA90R-H vs. WT-H, highlighting the core set of 24 shared genes and condition-specific transcriptional changes in early (JA90R) and late (L90R) stages of pollen development. (**c**) Heatmap showing the expression profiles of 24 shared DEGs across wild-type (WT) and HSP90 knockdown RNAi lines (L90R-C, L90R-H). The colour intensity represents the log2 fold-change values, with darker red indicating up-regulation and white to blue representing down-regulation.

In Panther Gene Class analysis of significantly down-regulated genes in L90R under normal conditions, DNA metabolism-related genes were enriched/overrepresented (Table S1). In L90R lines, we showed that there was a notable down-regulation of genes involved in protein folding, DNA metabolism, and DNA repair (Figure 5b, c, d). Key replication-related genes, including DNA Polymerase Alpha (POLA), DNA Polymerase Subunit B (POL2B), and DNA Polymerase Gamma Subunit 1 (POLGAMMA1), were significantly down-regulated (Figure 5d). These genes are essential for DNA synthesis and cell division, particularly during gametogenesis (Gumi and Bello 2024). This down-regulation of DNA metabolism helps explain the structural defects of MGU nuclei in pollen.

Taken together, genes associated with DNA metabolism highlight the distinct response in L90R line, emphasising the specific role of HSP90 in maintaining genomic stability during pollen development.

### ABA signalling pathway is activated under control conditions in L90R

Next to DNA metabolism, another functional group of genes showed interesting behaviour in the L90R background. Several genes associated with the abscisic acid (ABA) signalling pathway were significantly up-regulated under control conditions. These included *DEHYDRATION-RESPONSIVE ELEMENT BINDING PROTEIN 2A* (*DREB2A*), *LONG HYPOCOTYL IN FAR-RED 1* (*HFR1*), *MITOGEN-ACTIVATED PROTEIN KINASE KINASE KINASE 17, 18,* (*MAP3K17, 18*), *SCARECROW-LIKE PROTEIN 13* (*SCL13*), *DETOXIFICATION EFFLUX CARRIER 48* (*DTX48*), and *S-PHASE KINASE-ASSOCIATED PROTEIN 2B* (*SKP2B*) (log₂(Fold Change) and p-value shown in Figure 6e). Transcriptomic analysis further revealed that the MAP3K17/18-MKK3-MPK1/2/7/14 cascade, involved in ABA signalling, was up- regulated in L90R lines. Additionally, other key ABA biosynthesis and signalling genes such as *ZEAXANTHIN EPOXIDASE* (*ZEP*) (involved in ABA biosynthesis) and *HFR1* (linked to light signalling and oxidative stress responses) were detected among the DEGs (Figure 6b,e). We also detected *HFR1* among the DEGs in 35S:HSFA1b under heat stress (Figure S3, Supplementary data S4).

**Figure 6.**
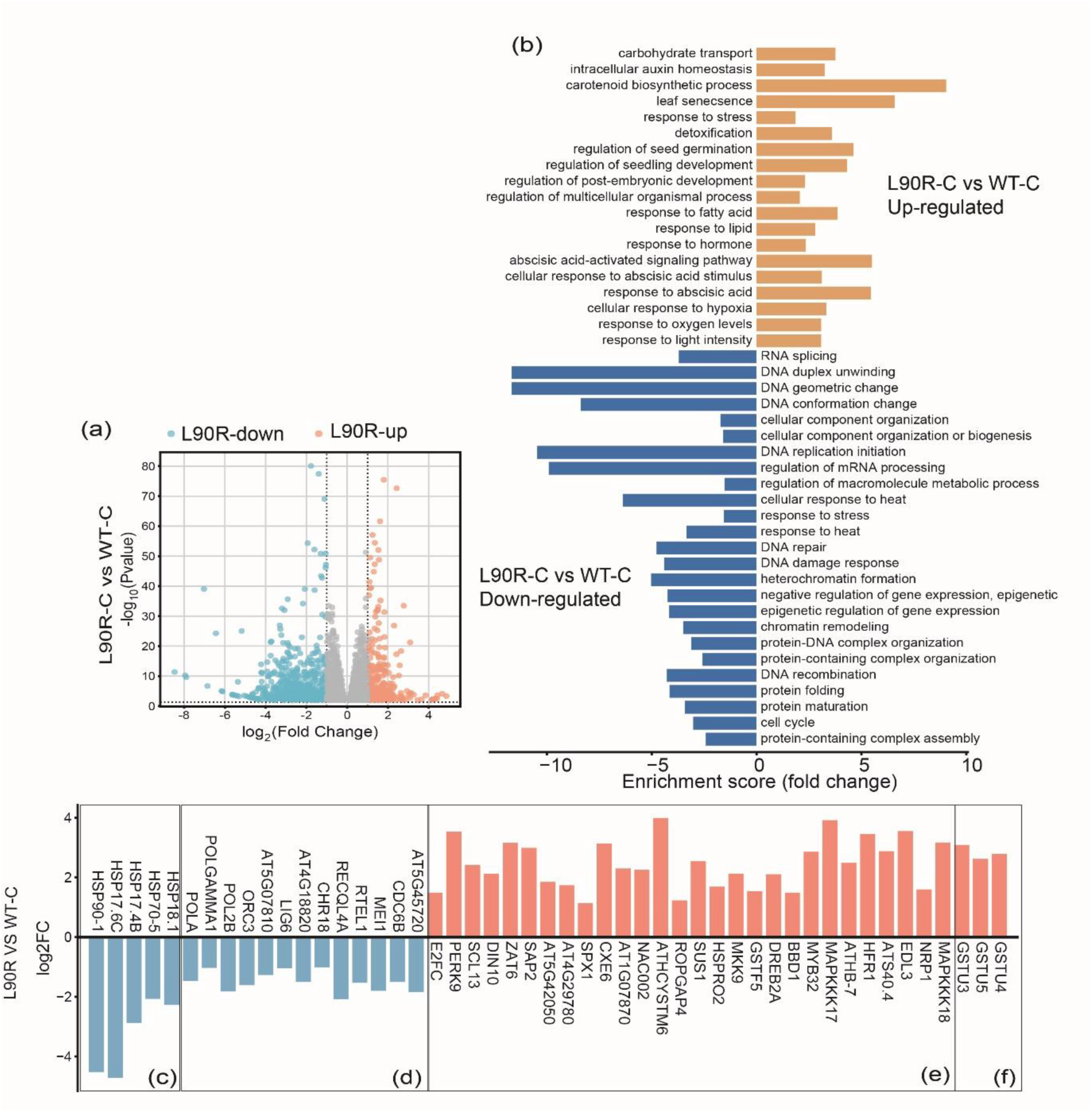
Differential gene expression analysis in pollen samples of WT and L90R line. (**a**) Volcano plot showing the distribution of gene expression changes (log₂ fold change) between WT and L90R. Genes with significant down-regulation are shown in blue, and genes with significant up-regulation are shown in orange. The x-axis represents the log₂ fold change, and the y-axis shows the -log₁₀(p-value), indicating the significance of the differential expression. (**c**) Bar graph showing up-regulated biological process (orange) and down-regulated biological process in L90R vs WT under control conditions **(c)** down-regulation of HSP90 and other HSPs in L90R **(d)** down-regulation of genes related to DNA metabolism **(e)** up-regulation of genes related to ABA response pathway **(f)** up-regulation of genes involved in ROS detoxification.

Glutathione S-Transferase family members (antioxidant enzymes for detoxification) including *GST4*, *GST3*, and *GST5*, were also highly up-regulated in L90R lines (Figure 6f). These genes encode antioxidant enzymes that reduce reactive oxygen species (ROS) build-up, particularly during anther development. In summary, several groups related to phytohormone signalling or stress metabolism showed up-regulation in L90R line.

### Heat response in *hsp90* RNAi lines shows distinct mechanisms in early and late pollen

In addition to previous results that highlighted the vital role of HSP90 in preserving genomic stability and developmental processes in pollen development, we further analysed the transcriptional differences between JA90R and L90R lines under heat stress (Figure 7a, b).

**Figure 7.**
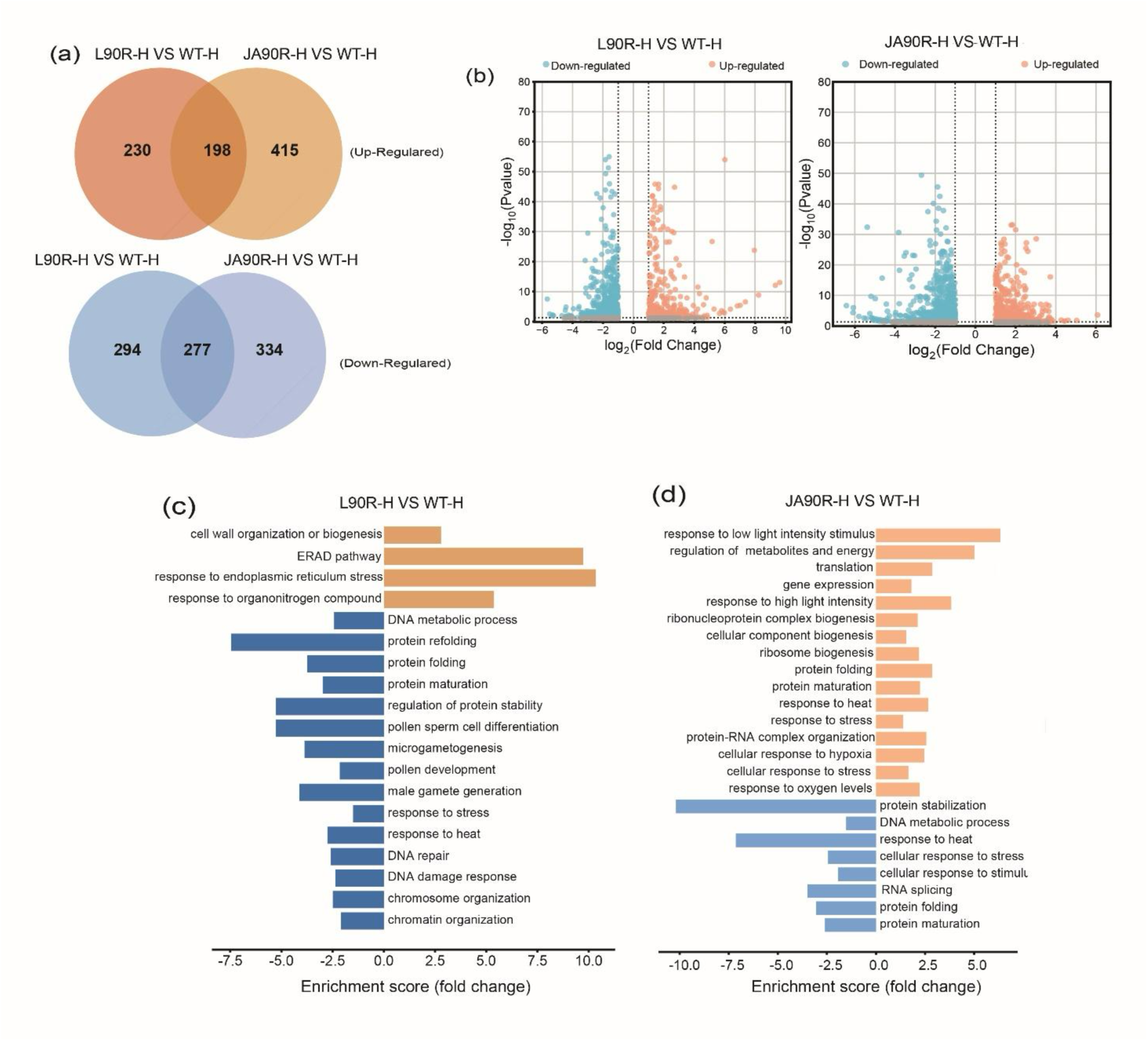
(a) Venn diagram showing the number of up- and down-regulated differentially expressed genes (DEGs) in mature pollen (DEGs) in JA90R vs WT, and L90R vs WT under heat stress (H) (37°C). Overlapping regions indicate common DEGs between the comparisons. **(b)** Volcano plots displaying the distribution of DEGs based on log₂(Fold Change) and -log₁₀(p-value) for WT vs L90R (left), and WT vs JA90R (right) under heat stress (H) (37°C). Points in orange and blue represent significantly up-regulated and down-regulated genes, respectively. **(c, d)** the biological processes represented by the up-regulated and down-regulated genes in **(c)** L90R and **(d)** JA90R under heat stress (H). Blue bars representing down- regulated biological processes and orange bars showing up-regulated processes.

In both RNAi lines, 198 genes are consistently up-regulated, representing similar heat stress response mechanism. On the other hand, 225 and 427 genes are exclusively increased in L90R and JA90R under heat stress, respectively. L90R-H has 294 uniquely down-regulated genes, while JA90R-H has 334 uniquely down-regulated genes, showing that both RNAi lines have distinct gene expression changes under heat stress, suggesting unique pathways affected during early and late pollen development (Figure 7a). The heatmap of the top 50 DEGs in L90R-H and JA90R-H lines under heat stress shows their distinct transcriptional responses to heat stress (Figure S4). In gene ontology (GO) enrichment analysis, L90R line exhibits a more pronounced down-regulation of genes associated with pollen-specific processes, DNA repair, and protein folding (Figure 7e), indicating disturbances in DNA and nucleus maintenance in pollen development. Among L90R up-regulated genes, significant enrichment of groups connected to the endoplasmic reticulum stress response, protein folding, and cell wall structure were detected (Figure 7d).

JA90R has an extensive range of up-regulated GO terms, with strong activation of metabolic and energy-related pathways, including ribosome biogenesis, translation, and oxidative stress response (Figure 7e). Down-regulation GO terms were less variable, containing pathways such as DNA metabolism and cellular stress responses. This indicates an emphasis on preserving cellular function and viability under stress and potential impairment in early pollen stages under stress conditions.

This analysis highlighted gene groups differentially expressed in both RNAi lines, we showed that JA90R line emphasizes overall stress response by increasing expression of metabolic and protein stability mechanisms, possibly alleviating significant developmental defects. L90R exhibits pronounced deficiencies in DNA stability and pollen-specific processes, resulting in compromised late-stage pollen development under heat stress. These results suggest that JA90R shifts more towards general cellular response, whereas L90R suffers from disruptions of key developmental pathways under heat stress.

### HSP90 maintains genome integrity in male gametophyte under heat stress

We will now turn to genes important for genomic maintenance and stability, further exploring the late-stage suppression of *HSP90* that leads to MGU defects in the L90R line. Among the genes, we detected the down- regulation of *ATP-DEPENDENT DNA HELICASE II 70 KDA SUBUNIT* (*KU70*), *DNA REPAIR PROTEIN RAD4*, *RAD5*, *RAD5B*, *MUTS HOMOLOG 5* (*MSH5*) and *MINICHROMOSOME MAINTENANCE COMPLEX COMPONENT 9* (*MCM9*) that are central to DNA repair under heat stress (Figure 8a). We also detected the up-regulation of *POSTMEIOTIC SEGREGATION INCREASED 1 HOMOLOG* (*PMS1*) and *CYCLIN-DEPENDENT KINASE A-1* (*CDKA-1*) in L90R with higher log_2_Fold change compared to WT under heat stress (Figure 8a). *PMS1* in plants, specifically in *A. thaliana*, is part of the MutLa complex, which plays a crucial role in the DNA mismatch repair (MMR) system. It is involved in correction of DNA biosynthetic errors, by which it maintains genomic stability (Spampinato, 2017).

**Figure 8.**
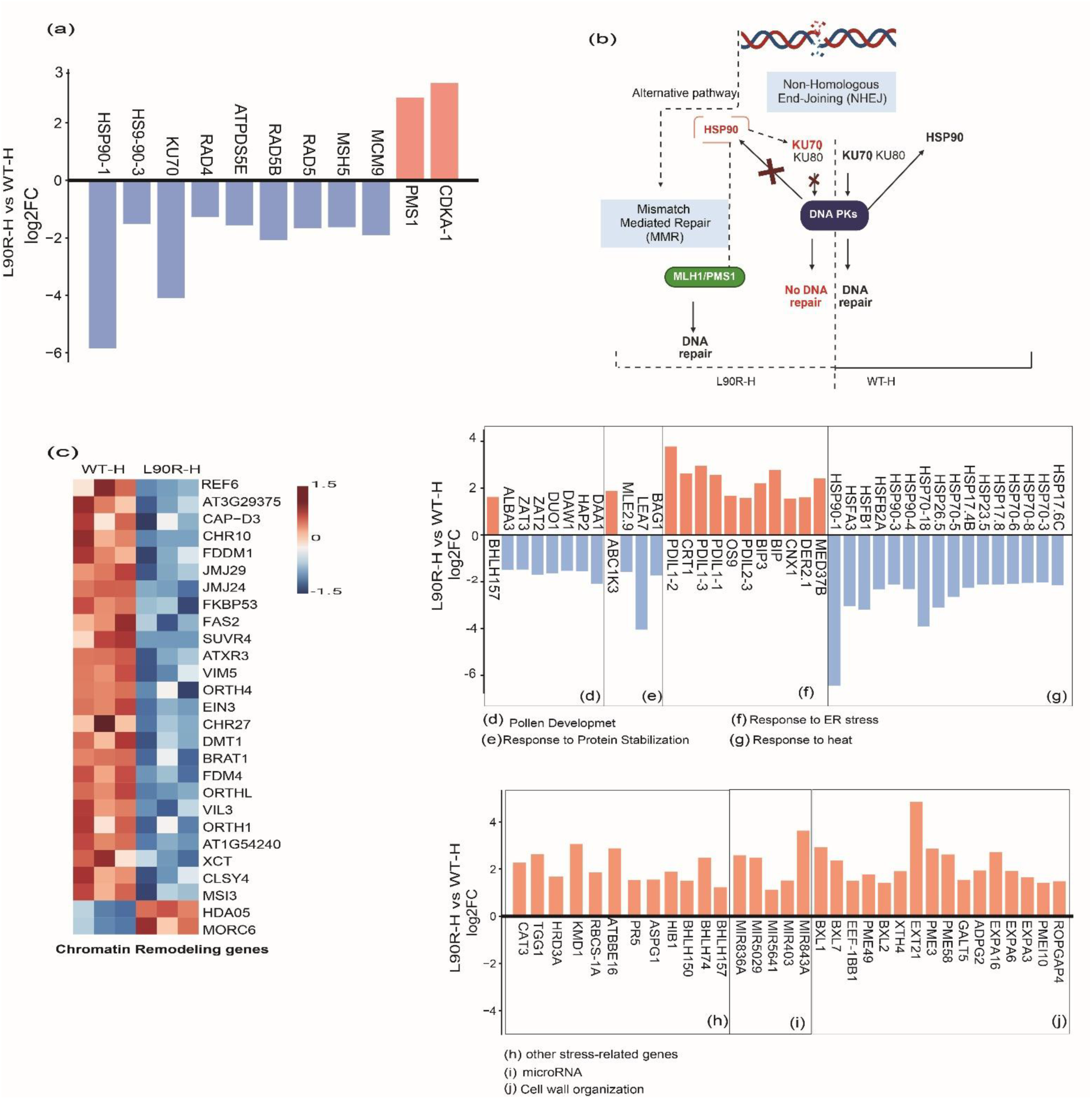
(a) log₂(Fold Change) values for selected genes involved in DNA repair in WT vs L90R under heat stress (H) (**b**) Model illustrating critical interactions between genes in DNA repair pathways, emphasizing the dependence of Non-Homologous End Joining (NHEJ) on HSP90 and the complementary function of Mismatch Mediated Repair (MMR) in L90R-H by up-regulating PMS1 **(c)** Heatmap of chromatin remodelling related genes in WT vs L90R under heat stress (H). (**d-j**) log₂(Fold Change) values for selected genes involved in stress response in WT vs L90R under heat stress. Blue bars represent up- regulated genes, while orange bars represent down-regulated genes.

In *HSP90* RNAi lines subjected to heat stress, there was a notable down-regulation of chromatin-associated and transcriptional regulatory genes (see heatmap on Figure 8c), pointing to a transition in chromatin-associated and transcriptional regulatory pathways to address stress and optimize cellular resource distribution. The genes associated with histone modification (*REF6*, *SUVR4*, *ATXR3*), chromatin remodelling (*CLSY4*, *XCT*), and transcription factor binding (*EIN3*) direct gene expression and epigenetic processes. L90R have compromised chaperone action, hindering protein structural integrity under heat stress to preserve resources and avert abnormal gene expression under unstable situations (Figure 8c).

The immense landscape of genomic balance, DNA structure and DNA repair and stability genes dysregulated in L90R line clearly show that nucleus maintenance is impaired, corresponding with the observed phenotype.

### Heat stress adaptation and structural modifications in L90R line

We show here the impact of HSP90 deficiency during the late stages of male gametophyte development on DNA repair systems and chromatin remodelling genes. Furthermore, we analysed the expression patterns of heat stress response-related genes in the L90R lines under heat stress conditions to identify the regulatory pathways involved in maintaining cell survival in the combined context of HSP90 deficiency and heat stress. Genes involved in heat responses including *HSFA3*, *HSFB1*, *HSFB2*, *HSP70* and *sHSPs* are down-regulated in L90R compared to WT under heat stress (Figure 8g). We showed a similar down-regulation of *HSFs* in JA90R, and L90R in Figure S5. The data showed that L90R exhibited a reduced expression of essential heat stress response genes and DNA repair genes, while selectively activated ER stress pathways, possibly as an adaptive response. The induced *PDIL1*-1, *PDIL1-2*, and *PDIL1-3* activates the Unfolded Protein Response (UPR) pathway, which ensures proper protein folding and protects against ER stress (Figure 8f). Cytoplasmic stress responses, principally mediated by HSP90 and HSFs, allow correct protein folding and proteostasis under stress (Fragkostefanakis et al., 2016). When cytoplasmic heat stress routes fail, ER stress responses, such as UPR activation via PDILs and BIPs, serve as a backup mechanism (Zhang & Kaufman, 2008). Several studies have found that ER stress-related pathways become especially important during late-stage pollen development, as proper membrane protein folding, secretion, and vesicular transport are required for pollen tube elongation and fertilization (Fragkostefanakis et al., 2016; Yang et al., 2017).

Genes like *ABC1-LIKE KINASE 3* (*ABC1K3*) associated with redox regulation show increased activity in L90R-H, suggesting stress-induced protein stabilization mechanisms (Figure 8e). In pollen developmental aspects, we showed the down-regulation of *DUO POLLEN 1* (*DUO1*) and *ALBA-DOMAIN CONTAINING PROTEIN 3* (*ALBA3*), whereas the up-regulation of *BASIC HELIX-LOOP-HELIX PROTEIN 157* (*BHLH157*) suggests an adaptive response in L90R under heat stress (Figure 8d). *BHLH157* potentially controls downstream genes allowing the cell to adapt to stressors such oxidative damage or heat stress.

Other stress-responsive genes, including *CATALASE 3 (CAT3)*, *THIOGLUCOSIDE GLUCOHYDROLASE 1 (TGG1)*, and *KELCH REPEAT F-BOX PROTEIN 1 (KMD1)*, were up-regulated in the L90R line under heat stress compared to WT, suggesting their involvement in oxidative stress mitigation and detoxification processes (Figure 8h).*CAT3* encodes a catalase enzyme that detoxifies reactive oxygen species (ROS), while *GLUTATHIONE S-TRANSFERASE TAU 19 (GSTU19)* encodes a glutathione S-transferase involved in cellular detoxification, both highlighting key adaptation mechanisms under HSP90 deficiency. Among other stress-related genes, *HBI1 (HOMEOBOX-INTERACTING BASIC HELIX-LOOP-HELIX PROTEIN 1)* was also strongly induced. As a bHLH transcription factor family, *HBI1* plays a central role in maintaining cellular homeostasis by activating genes involved in oxidative stress tolerance and repair pathways (Figure 8h).Additionally, *HBI1* was detected among the DEGs in the 35S:HSFA1b transgenic line under heat stress conditions (Figure S3; Supplementary Data S4).

The up-regulation of genes associated with cell wall organization like *BETA-XYLOSIDASE* 1 (*BXL1*), *EXTENSIN 21* (*EXT21*), and *XYLOGLUCAN ENDOTRANSGLUCOSYLASE/HYDROLASE 4* (*XTH4*) in L90R lines specifies that the cells are experiencing a structural and stress-related adaptation to preserve cellular integrity under adverse conditions (Figure 8i). Among up-regulated DEGs, we detected MicroRNAs highly up-regulated in L90R under heat stress (Figure 8j). MicroRNAs are responsible for silencing stress-sensitive genes, which protects the cell from maladaptive reactions. The participation of certain microRNAs is consistent with previous studies indicating their significance in post-transcriptional regulation under heat stress (Bokszczanin et al., 2013).

This multifaceted response underscores the complex regulatory networks that are modulated in hsp90 RNAi lines to cope with heat stress, involving both transcriptional and post-transcriptional mechanisms, as well as structural modifications.

## Discussion

One of the most vulnerable phases in angiosperm plant life cycle is the sexual reproduction. Successful reproduction of most angiosperm plant species depends on effective fertilization (Resentini, 2023), but also on the thermotolerance of pollen (Lohani et al., 2020; Fragkostefanakis et al., 2014; Mesihovic et al., 2016). Our study investigated how elevated temperatures influence gene expression across five distinct pollen developmental stages in *A. thaliana*: UN, EB, BC, TC and MP).

Upon exposing plants to heat stress (in our study represented by 37 °C for three hours), we observed a substantial increase in induced DEGs at the BC stage, exceeding 4,000, compared to 271–879 DEGs in other stages. This suggests that the BC stage is particularly sensitive to heat stress, undergoing extensive transcriptional reprogramming. Notably, the majority of DEGs were stage-specific, indicating unique responses to heat stress at each developmental stages. The unique transcriptomic profile of the stress response can be also linked to the dramatic shifts in the overall transcriptomic profile of the developmental stages in *A. thaliana* (Klodová et al., 2023; Sze et al., 2024).

Gene Ontology analysis revealed that "response to heat" and "protein (re)folding" were enriched across all stages, which agrees with a partial activation of general heat shock responses. However, other stress-related categories, such as "cellular response to hypoxia" were enriched only in TC and MP stages, suggesting specific aspects of stress adaptations. The BC stage exhibited unique enrichments in pathways related to RNA processing, organellar transcription, and metabolic processes, reflecting a broad transcriptional adjustment to heat stress.

Interestingly, among the heat-induced genes, only 23 were consistently up-regulated across all developmental stages, with 13 of these encoding sHSPs. The induction of sHSPs was more prominent in later stages (TC and MP) compared to early stages, indicating a developmental modulation in the amplitude of heat stress responses. These transcriptomic observations were supported also by our promoter motif analysis (Nevosád et al., 2025) that revealed that enrichment of the HSE in the promoters of highly transcribed genes in plants that underwent heat stress. The stage-specific HSE pattern thus emphasizes a coordinated regulatory architecture that supports heat response as pollen matures. This pattern contrasts starkly with other cis-elements like the *telo*-box motif, enriched at BC under heat but absent in MP. The *telo*-box motif commonly found in promoters of ribosomal protein genes that are enriched in early pollen development (Schrumpfova et al., 2016; Klodová et al., 2022), underscoring a functional shift in regulatory strategies as development progresses. The TATA-box, previously linked to both general environmental responses and stress-responsive transcription (Pavlu et al., 2024), showed inconsistent enrichment. Similarly, the dehydration-responsive DRE/CRT motif (Agarwal et al., 2017) and the LAT52 motif, associated with late pollen development (Bate and Twell, 1998), were notably unresponsive across all developmental stages under heat stress. These findings suggest a limited role for these elements in the heat stress adaptation of the male gametophyte. Collectively, these results show a finely tuned, stage-dependent orchestration of heat stress responses, where both transcriptomic and promoter motif landscapes adapt dynamically, enabling specific stress resilience strategies across pollen development.

To further reveal the stage-specific impacts of *HSP90* down-regulation in male gametophyte, we generated RNAi lines targeting *HSP90-3* and *HSP90-1* at specific developmental stages in *A. thaliana*. For early-stage male gametophyte development, we used the *JASON* promoter, which is active during male meiosis and plays a crucial role in maintaining organelle positioning and ensuring proper chromosome segregation to drive *HSP90-3* and *HSP90-1* knockdown, creating the JA90R line. In late-stage pollen development, the *LAT52* promoter, predominantly active in the vegetative cell during pollen maturation, directed gene expression in the later stages of pollen development (Bate & Twell, 1998; Wielopolska et al, 2005; Storme & Geelen, 2013; Cabout et al., 2017).

The mature stage of male gametophyte development is marked by rapid cell division, chromatin remodelling, and sperm cell differentiation. It requires meticulous coordination of DNA replication and repair processes (Borg & Berger, 2015). In our observation of late-stage L90R lines, we showed high percentage of pollen defects, particularly an impaired male germ unit organisation and nucleus structure under heat stress conditions. Using transcriptomic analysis, we linked this impaired MGU phenotype to the down-regulation of DNA maintenance and metabolism genes in the L90R lines. The observed transcriptomic changes suggest that *HSP90* deficiency disrupts critical processes, such as DNA replication and chromosomal integrity, during late-stage gametogenesis. It is highly interesting how such a change of DNA regulating genes could cause massive observable changes to the nucleus structure.

Among the key DNA replication-related genes that play pivotal roles in maintaining genome integrity, we detected down-regulation of *POLA* (encoding DNA Polymerase I), *POL2B*, and *POLGAMMA1*. DNA Polymerase I (Pol I) is essential for chromosomal replication, maturing Okazaki fragments on the lagging strand (Makiela-Dzbenska et al., 2009). PolB fills gaps and displaces downstream Okazaki fragments. *POLGAMMA1*, the catalytic subunit of mitochondrial DNA polymerase, is crucial for replication and stability (Cupp and Nielsen, 2013). Mutations in POLGAMMA1 impair DNA binding and synthesis, leading to stalled replication and mitochondrial dysfunction. This dysfunction can cause oxidative stress, ATP depletion, and altered metabolic states (Silva-Pinheiro et al., 2021). Although we did not see a clear disruption in the cell division, which could be caused by replication defects, these genes can nevertheless be part of broader *HSP90*-mediated regulation of transcription. Moreover, the genes involved in DNA repair, chromatin remodelling, and DNA methylation were found to be differentially expressed in the L90R lines compared to the wild-type under heat stress (Figure 9a,b,c). This suggests that the loss of HSP90 function impairs the cells’ ability to maintain genomic integrity and adapt to heat stress, leading to the impaired MGU phenotype defects. HSP90 interacts with proteins across multiple DNA repair pathways, including members of the Non-Homologous End Joining (NHEJ) and Homologous Recombination (HR) (Dubrez et al., 2020). HSP90 was shown to stabilize DNA-PKcs, a core component of NHEJ, and facilitates the recruitment of repair machinery to sites of DNA damage. HSP90 also, ensures proper folding and localization of BRCA1/2 and Rad51, two critical HR proteins, which is essential for accurate repair of double-strand breaks (Dubrez et al., 2020). Importantly, *KU70*, a critical component of the NHEJ pathway, was found to be strongly down- regulated in the L90R lines under heat stress (Figure 8b), reinforcing the idea that compromised DNA repair capacity contributes to the observed reproductive defects.

**Figure 9.**
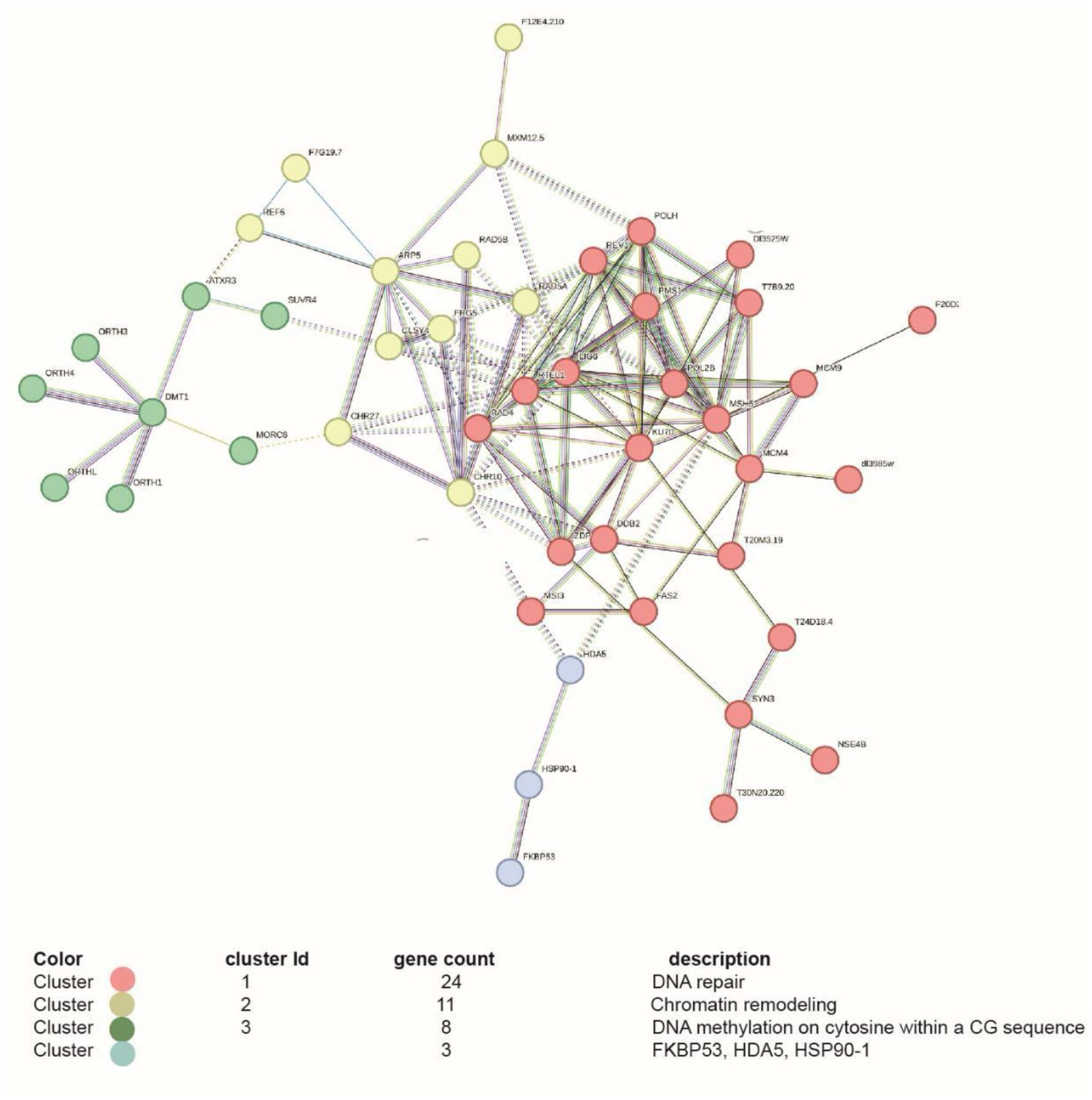
The STRING network analysis illustrates the protein-protein interaction (PPI) network of key DEGs in L90R under heat stress related to DNA Repair, Chromatin remodelling and DNA methylation. Each cluster is shown by different colour indicated in the table.

Under heat stress, L90R exhibits an increased expression of genes associated with DNA repair and DNA damage response, suggesting a potential role in maintaining genetic stability through the up-regulation of these biological processes (Figure 8a, b).Surprisingly, we detected the up-regulation of *PMS1* in L90R compared to WT under heat stress (Figure 8a). *PMS1* in plants, specifically *A. thaliana*, is part of the MutLa complex, which plays a crucial role in the DNA mismatch repair (MMR) system. It is involved in correcting DNA biosynthetic errors, thereby maintaining genomic stability (Spampinato, 2017). The up-regulation of this gene contradicts the usual trend of DNA maintenance down-regulation seen in L90R. A new regulatory mechanism in certain gene groups that bypassed typical HSP-related gene expression regulation might help to explain it.

Low levels of HSP90 diminish its chaperone actions and this protein structural integrity under heat stress. In these conditions, cells likely down-regulate certain chromatin and regulatory genes to preserve resources and avert abnormal gene expression under unstable situations (Leach et al. 2016). We observed this mechanism in L90R by detecting down-regulated genes associated with histone modification (REF6, *SUVR4*, *ATXR3*), chromatin remodelling (*CLSY4*, *XCT*), and transcription factor binding (*EIN3*), all involved in direct gene expression and epigenetic processes. This down-regulation may have a dual function. The deficiency of HSP90 increases transcriptional errors and genomic instability. Hence, diminishing these chromatin regulators may mitigate the misexpression of stress-sensitive genes. This deliberate biological modification restricts the capacity of chromatin and transcriptional machinery to stabilize the genome when HSP90-dependent stabilization pathways are less effective. This supports the model of canalization provided by HSP90—traditionally characterized at the protein folding level—which we now observe reflected at the transcriptional level as well. This interpretation aligns with the growing understanding of HSP90’s role in maintaining both genetic and epigenetic robustness under stress (Zabinsky et al., 2019; Geiler-Samerotte et al., 2019).

In parallel, adaptive alterations may emphasize essential stress-response pathways and prevent uncontrolled gene expression during heat stress in the compromised *HSP90* RNAi line context. On the other hand, *MORC6*, a gene associated with heterochromatin formation and transposable element silencing, is likely up- regulated in L90R lines during heat stress to improve genome stability by suppressing unwanted transcriptional activity and transposable element mobilization, which can be harmful during stress (Brabbs et al., 2013). The expression of *HDA05*, a histone deacetylase, is also increased, facilitating chromatin compaction and gene repression in order to save energy and avoid incorrect gene expression (Luo et al., 2017). This combined up-regulation of *MORC6* and *HDA05* implies a compensatory response in L90R lines, maintaining chromatin structure and genomic stability even when cellular conditions are disrupted by the mutation and heat stress. Supporting this model, interaction analyses of DEGs in heat-stressed L90R lines revealed coordinated activity between DNA repair, chromatin remodelling, and DNA methylation pathways, as illustrated in Figure 9.

In addition to the partially compromised DNA repair mechanisms observed in the L90R line, a distinct and substantial group of dysregulated genes were those associated with the abscisic acid (ABA) signaling pathway, which were up-regulated in L90R even under control conditions (Figure 6c, g). The genes linked with the up-regulation of the ABA signalling pathway include *DREB2A*, *HFR1*, *MAP3K17*, *MAP3K18*, *SCL13*, *DTX48*, and *SKP2B* (Figure 6). *MAP3K17* is identified as an ABA-inducible mitogen-activated protein kinase (MAP3K) that plays a role in ABA signalling. It is part of the MAP3K17/18-MKK3- MPK1/2/7/14 cascade, which is up-regulated in L90R under control conditions that were implicated in regulating various aspects of plant growth and stress responses (Matsuoka et al., 2018).

Further, HSFA1s can trigger HS-inducible gene expression via transactivation of other HSFs such as *DREB2A*, *HSFA2s*, *HSFA3s*, *HSFA7s*, *HSFBs*, and others (Ohama et al., 2017: Jacob et al., 2017). The HSFA1 regulates the induction of several other HSFs, including *HSFA2*, *DREB2A,* HSFA7s, and even certain micro-RNAs that initiate the expression of HSPs and other heat stress-inducible genes, which eventually orchestrate physiological responses precipitating thermotolerance acquisition (Zenda et al, 2022). Among these, *DREB2A* is critical for abiotic stress tolerance, particularly drought and high salinity, in addition to its role in the HS response (Mizoi et al., 2019).

*HFR1*, *ZEP*, *SCL13*, *DTX48*, and *SKP2B* indicate together a broader adaptive strategy since *HFR1* modulates light signalling and oxidative stress response (Jing and Lin 2020), while *ZEP* primes for ABA production, crucial for drought resilience, by catalysing a key step in abscisic acid biosynthesis (Long et al., 2019). *HFR1* was also identified as a differentially expressed gene in 35S:HSFA1b under heat stress conditions (Albihlal et al., 2018) (Figure S3, Supplementary data S4). HSFA1b, a master regulator of heat stress responses, controls the transcription of genes involved in various protective mechanisms. The detection of *HFR1* as a differentially expressed gene implies it could be a downstream target or part of the regulatory network governed by HSFA1b. *HFR1* may contribute to fine-tuning stress signalling pathways, such as those associated with ABA (abscisic acid) signalling, and influence growth and developmental processes under thermal stress (Bulgakov et al., 2022; Albihlal et al., 2018). *DTX48* aids detoxification as part of a transporter family involved in managing xenobiotics and stress-related compounds (Lu et al., 2019). *SKP2B* helps in targeted protein degradation by facilitating ubiquitin-mediated degradation, which is essential for clearing misfolded proteins and maintaining cellular stability (Del Pozo and Manzano, 2014).) (Figure 6c). Some studies suggest that HSFs can influence transcription factors involved in broader stress responses, including the drought-responsive *DREB2A*, especially when cellular stress overlaps (e.g. heat combined with drought) (Schramm et al., 2008; Yoshida et al., 2008; Sakuma et al., 2006). HSFs-targeted genes continue to play an important role in controlling the expression of protective genes. The increased expression of *DREB2A* and other transcription factors (such as *HBI1* and *bHLH50*) triggers the transcription of genes involved in adaptation to stress (Zenda et al, 2022). While *DREB2A* is a target of HsfA3, it is also part of a broader network of transcription factors and regulatory mechanisms that enable plants to adapt to various environmental stresses (Sato et all 2014). HBI1 is known to control ROS homeostasis and influence gene expression related to oxidative stress and heat tolerance (Prodromou, 2016). The involvement of HSF1 in stress granules, as well as its interactions with transcriptional activators like c-MYC, suggests that it has the capacity to impact gene expression broadly, perhaps regulating DNA repair responses indirectly. HSP90 sequesters HSF1 in the cytoplasm, suppressing its activity. Upon HSP90 inhibition, HSF1 is released, potentially driving oncogenic processes, which may indirectly influence DNA repair mechanisms (Xu, et al., 2023).

At last, genes related to detoxification like *GST4*, *GST3* and *GST5* were detected up-regulated in L90R line (Figure 6h). GSTs play a crucial role in plant stress responses by scavenging ROS and detoxifying harmful metabolites generated during heat stress. Previous studies have shown that GST genes are often co-expressed with HSFs under stress conditions, indicating their involvement in the heat stress response and cellular protection mechanisms. For example, HSF-dependent transcriptional regulation of GSTs has been documented in several plant species, such as Arabidopsis and wheat, where GSTs contribute to enhanced stress resilience by reducing oxidative damage and maintaining cellular homeostasis. In *A. thaliana*, HSFA1 and HSFA2 transcription factors have been found to regulate GST expression, further supporting their role in heat-induced detoxification mechanisms (Nishizawa et al., 2006).

In summary, our study offers understanding of the critical sensitivity of pollen development to heat stress, particularly at BC stage when we found notable transcriptional reprogramming. The stage-specific enrichment of heat stress response pathways emphasizes the complexity and adaptation of male gametophyte development under increased temperatures. The essential role of chaperones in preserving protein stability and cellular function under heat stress is shown by the constant up-regulation of sHSPs and developmental regulation of HSP70 and HSP90 expression. Under heat stress, our results show the critical role of HSP90 in preserving genomic integrity and cellular functions during the late stages of male gametophyte development (Figure 10). Showing the dependence of these processes on the expression of HSP90 chaperons, the observed transcriptome changes in the L90R line show crucial defects in DNA repair mechanisms, chromatin remodelling, and stress response systems. Added to that, up-regulation of important stress-response genes, such as those involved in antioxidant activity and transcriptional regulation, suggests a compensatory mechanism to mitigate the impacts of HSP90 deficiency. Cytoplasmic stress responses, such as UPR activation via PDILs and BIPs, serve as a backup mechanism for proper protein folding in HSP90 deficiency under heat stress. Down-regulation of chromatin-related genes in HSP90-deficient conditions could have served a dual role. The decrease of HSP90 first increases genomic instability, then reducing the expression of chromatin regulators in response could allow to avoid the misexpression of stress-sensitive genes, therefore preserving genome integrity. These results show the complex interaction among heat stress, HSP90 function, and male gametophyte viability, therefore providing understanding of the molecular adaptations required for successful reproduction under heat stress.

**Figure 10.**
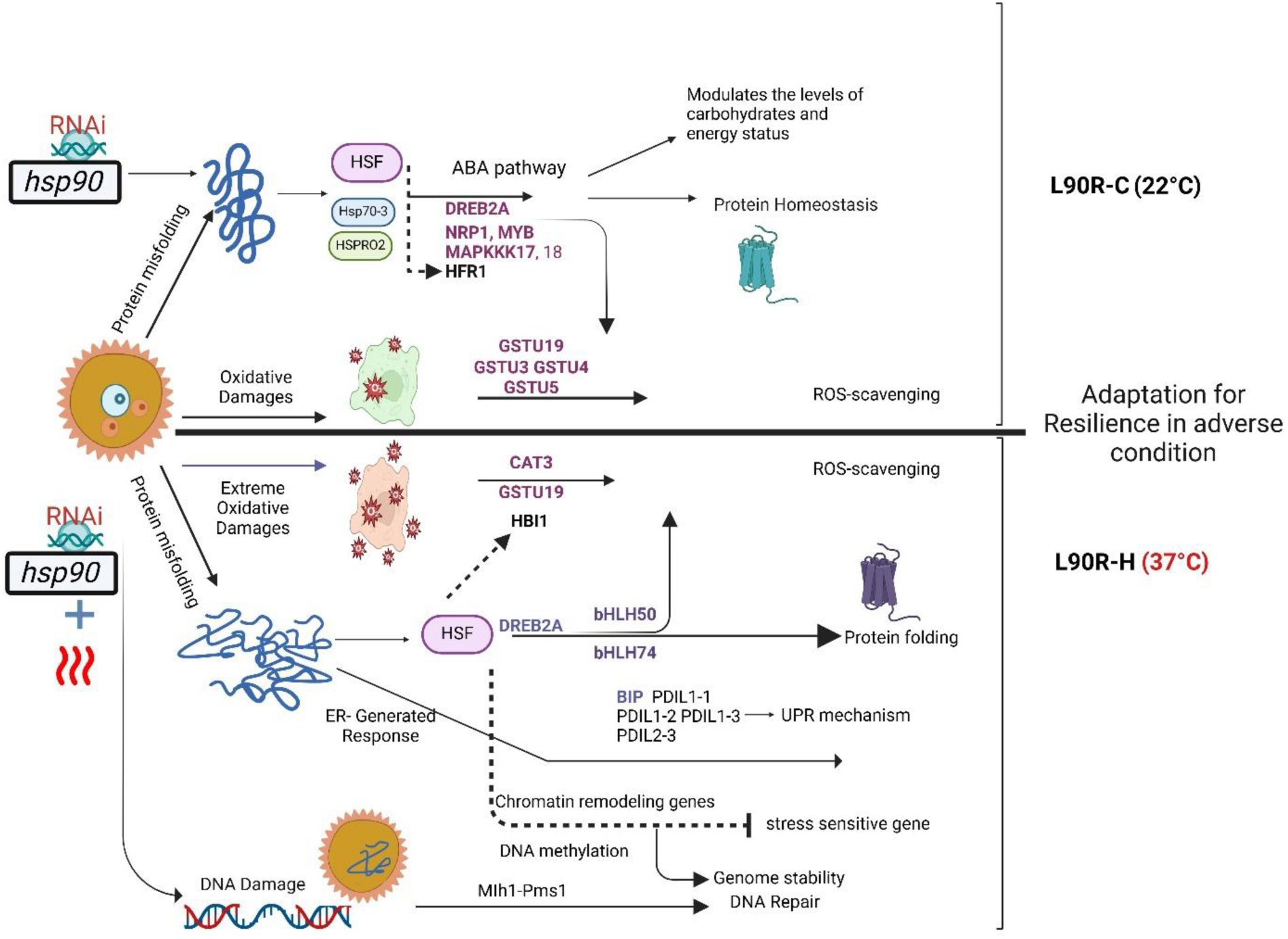
This model highlights the regulatory modifications in *HSP90* RNAi lines that facilitate resilience under adverse conditions via a coordinated network of protein folding, reactive oxygen species scavenging, and stress signalling pathways. L90R under normal conditions (22 °C) – L90R-C. In control conditions, HSF-targeted genes are involved in regulating the ABA pathway, influencing carbohydrate and energy levels to maintain homeostasis. Target genes like *DREB2A*, *NRP1*, *MYB* aid in reactive oxygen species (ROS) scavenging through *GSTU3*, *GSTU4*, *GSTU5* and *GSTU19* enzymes, which mitigate oxidative damage. (**b**) L90R under heat stress (37 °C) – L90R-H: *HSP90* knockdown impaired oxidative damage, leading to shift in stress response regulation by initiating ER (endoplasmic reticulum)-generated stress responses, including activation of *DREB2A* and HSF, which regulate genes like *bHLH50* and *bHLH74* for enhanced protein folding and stress resilience. ROS scavenging is intensified with the up-regulation of *CAT* and *GSTU19*. The unfolded protein response (UPR) mechanism is activated, with proteins such as BIP and various PDIL proteins promoting cellular stability. Chromatin Remodelling genes and DNA methylation related aid in silencing sensitive genes and increasing the genome stability. DNA repair by up-regulation of Pms1 complex part of the mismatch repair (MMR) pathway contributing to genome stability and the stress adaptation.

## Conclusion

With a focus on the function of HSP90s in regulating heat stress response, this study provides a comprehensive understanding of the transcriptional responses to heat stress in male gametophyte development. Our results reveal a stage-specific heat stress response in pollen development wherein most differently expressed genes unique to each stage with highest transcriptional changes in BC stage, while the strongest enrichment of HSEs found in mature pollen. This focuses on how dynamically stress responses are regulated during male gametophytes development, influenced by HSPs levels and cellular stress adaptation mechanisms.

The stage-specific effects of HSP90 knockdown show its critical role in maintaining cellular stability and genomic integrity in the late-stage development of male gametophytes under heat stress. Together with to its known function as a molecular chaperone, HSP90 regulates critical stress responses, including the regulation of heat shock proteins, the activation of the unfolded protein response, and DNA repair mechanisms, therefore preserving genomic integrity. Its deficiency disrupts redox homeostasis, alters cell wall composition, and triggers stress-induced cellular modifications. These results provide important insights into the molecular processes behind heat stress tolerance in pollen, creating a foundation for targeted breeding techniques aimed at improving crop stability and yield in response to climate change.

## Material and Methods

### Plant material, growth conditions and heat stress treatment

*Arabidopsis thaliana* Col-0 plants were grown in a controlled environment with 16-hour light/8-hour dark cycles at 22°C and 50% relative humidity. Seeds were sown in ½ MS medium (COMPANY) and were stratified for two days at 4°C before in vitro germination at conditions listed above. Then, 7-days old seedlings were transferred to a growth chamber under the described conditions and cultivated for 6–8 weeks until inflorescences fully developed.

For heat stress treatment, we exposed *A. thaliana* plants at the inflorescence stage to a temperature of 37°C for three hours to induce heat stress. Control plants were maintained at 22°C under identical light and humidity conditions. After 24-hour recovery phase, inflorescences were harvested from both the heat-treated and control plants. These were then processed to isolate pollen at distinct developmental stages using the Percoll gradient separation protocol (see below) based on the protocol by Dupľáková et al. (2016).

### Assembly of RNAi expression cassettes and transgenic line selection

To target the expression of HSP90-3 and HSP90-1 during distinct developmental stages, a 602 bp fragment of the HSP90-3 sequence—selected for its high sequence similarity to HSP90.1—was amplified using gene-specific primers and cloned into the GoldenBraid 3.0 system in both sense and antisense orientations to generate a hairpin RNA structure for RNAi. To enhance the efficiency of gene silencing, an intron derived from the pHANNIBAL vector was inserted between the sense and antisense sequences, ensuring proper processing of the hairpin RNA.

Promoters were strategically selected to regulate RNAi induction at specific developmental stages. The GoldenBraid—domesticated JASON promoter (AT1G06660) was used to drive RNAi expression during early pollen development, while the LAT52 promoter was employed for later stages (Wielopolska et al, 2005; Bate & Twell, 1998; Storme & Geelen, 2011). Both RNAi expression cassettes are terminated by NosT terminator (Bevan, 1984). Primers used for 90R domestication or elsewhere in the cloning are listed in supplementary table S2.

Each RNAi expression cassette was ligated with KanFAST selection cassette into Omega2 level binary vector and transformed into *A. thaliana* (Col-0) plants through Agrobacterium tumefaciens-mediated transformation. Floral dipping was done according to (Clough et al, 1998). Seed harvested from the T_0_ transformed plants were selected on ½ MS media with 100 mg/L Kanamycin used as a selection, otherwise plants were cultivated under standard conditions. Independent transgenic lines were further cultivated and selected until a uniform homozygous line in T_3_ generation, which were used for the analysis.

### Separation of developmental stages using a Percoll gradient

Following the method by Dupľáková et al. (2016), our protocol for separating *A. thaliana* pollen developmental stages relied on a stepwise Percoll gradient to isolate specific pollen stages. First, inflorescences were collected, and the pollen was gently released by homogenizing the floral tissue in a chilled 0.1 M d-mannitol. Prior to loading the pollen suspension onto the Percoll gradient, a one-step cleanup modification was introduced to remove debris and improve the purity of the sample. This step involved centrifuging the suspension to eliminate floral tissue fragments, ensuring a cleaner starting material for gradient separation. The pollen suspension was subjected to a centrifugation process using a discontinuous Percoll gradient with multiple layers of increasing Percoll density. This setup allowed pollen grains to settle at distinct layers based on their developmental stage and density. First Percoll gradient (75%/65%/20%/10%) enabled separation into uni-cellular, bi-cellular, and tri-cellular fractions, while a subsequent gradient (55%/45%/35%) refined these stages further, ensuring up to 80-90% purity.

### Phenotype analysis

To describe pollen morphology and anatomy, 4′,6-Diamidino-2-phenylindole (DAPI) staining was used (Vergne et al., 1987). The screening of mature pollen samples was conducted using Zeiss Axiovert 200M under Bright-field microscopy and UV light that excites the DAPI (LSM200). We scored our screening for pollen phenotype defects following the approach described here (Reňák et al., 2012).

### Pollen thermotolerance assay and in vitro pollen germination

We conducted a pollen thermotolerance assay to evaluate the impact of heat stress on pollen viability and germination. This assay involved subjecting flower buds to a controlled heat stress treatment at 37°C for a duration of three hours. Following this heat exposure, the buds were allowed to recover under normal conditions in 37°C for 24 hours. Subsequently, the pollen collected from these buds was analyzed for two key parameters: viability and germination. We followed the method described by M. P. Alexander (1969) for testing pollen viability. Boavida and McCormick’s protocol (2007) method for in vitro pollen tube growth was followed. The pollen germination rate was determined after 8 h of *in vitro* growth.

### RNA extraction

Total RNA was isolated from populations of isolated microspores or developing pollen (UNM, BCP, TCP and MPG). The RNeasy Mini Kit (Qiagen, Valencia, CA, USA) was used to extract total RNA from three biological replicates of isolated pollen stages. Isolated RNA was treated with RQ1 RNAse-free DNase I treatment, (Promega in Maryland, MD, United States). The NanoDrop One instrument (Thermo Fisher Scientific, Waltham, MA, USA) was used to measure the RNA quantity and quality. The RNA quality was also verified by electrophoresis in a 2% agarose gel.

### RNA-seq data processing, mapping and assembly of reads

The quality of single-end raw readings was assessed using FastQC version 0.11.8 (Wingett and Andrews 2018) and Cutadapt version 1.9.1 (Martin 2011). The quality reads (phred score > 20) were stripped of technical sequences using Cutadapt software and then aligned to the *A. thaliana* reference genome (version TAIR10) obtained from Araport (Pasha et al. 2020) using STAR version 206.1a (Dobin et al. 2013). The GTF annotation file acquired from Araport was used for STAR index construction. Gene counts, together with adjusted TPM levels, were obtained using RSEM (Li and Dewey 2011). The data were imported into RStudio using Tximport (Soneson et al. 2015) and further processed. For differential expression analysis, DESeq2 version 3.8 (Love et al. 2014) was used, using adjusted p-values as the statistically significant thresholds in both studies.

### Data visualization and analysis thresholds

The lists of DEGs were annotated with gene names and symbols derived from ThaleMine v5.0.2 (Pasha et al. 2020). GO Enrichment for biological processes was analysed by Panther19.0 with Fisher’s Exact test with False discovery rate (FDR) correction. Principal Component Analysis (PCA) was performed based on the top 1000 differentially expressed genes (DEGs). STRING network analysis was applied to DEGs with >1.5-fold change and p-value <0.01. HSE motifs were analyzed using the GOLEM tool (Nevosád et al., 2025; https://golem-dev.ncbr.muni.cz/).

## Acknowledgements

The authors thank RECROP COST Action (www.cost.eu) CA22157 for networking opportunities. This work was supported by the Czech Science Foundation projects 21-15841S (P.P.S.), 23-07000S and 24- 10653S (D.H.), and by the ERDF Programme Johannes Amos Comenius project TowArds Next GENeration Crops, reg. no. CZ.02.01.01/00/22_008/0004581. The authors gratefully acknowledge the financial support from the Mobility Plus Program with DAAD (DAAD-23-06). The bioinformatic analyses performed by P.P.S. and J.R. were further supported by the Ministry of Education, Youth and Sports of the Czech Republic – project INTER-COST-LUC24 (LUC24056). We acknowledge the Imaging Facility of the Institute of Experimental Botany AS CR supported by the MEYS CR (LM2023050 Czech-BioImaging) and IEB CAS.

## Author contribution

CM, DH, and ZK designed the study. CM, ZK, KR performed the experiments and ZK and SF wrote the manuscript. BK, ZK, SF and CM analysed the RNA-seq data. PPS and JR analysed the data in GOLEM. CM, KR, JF, BK, SF, DH and PPS edited the manuscript. All authors approved the final manuscript.

---

**Figure S1.**
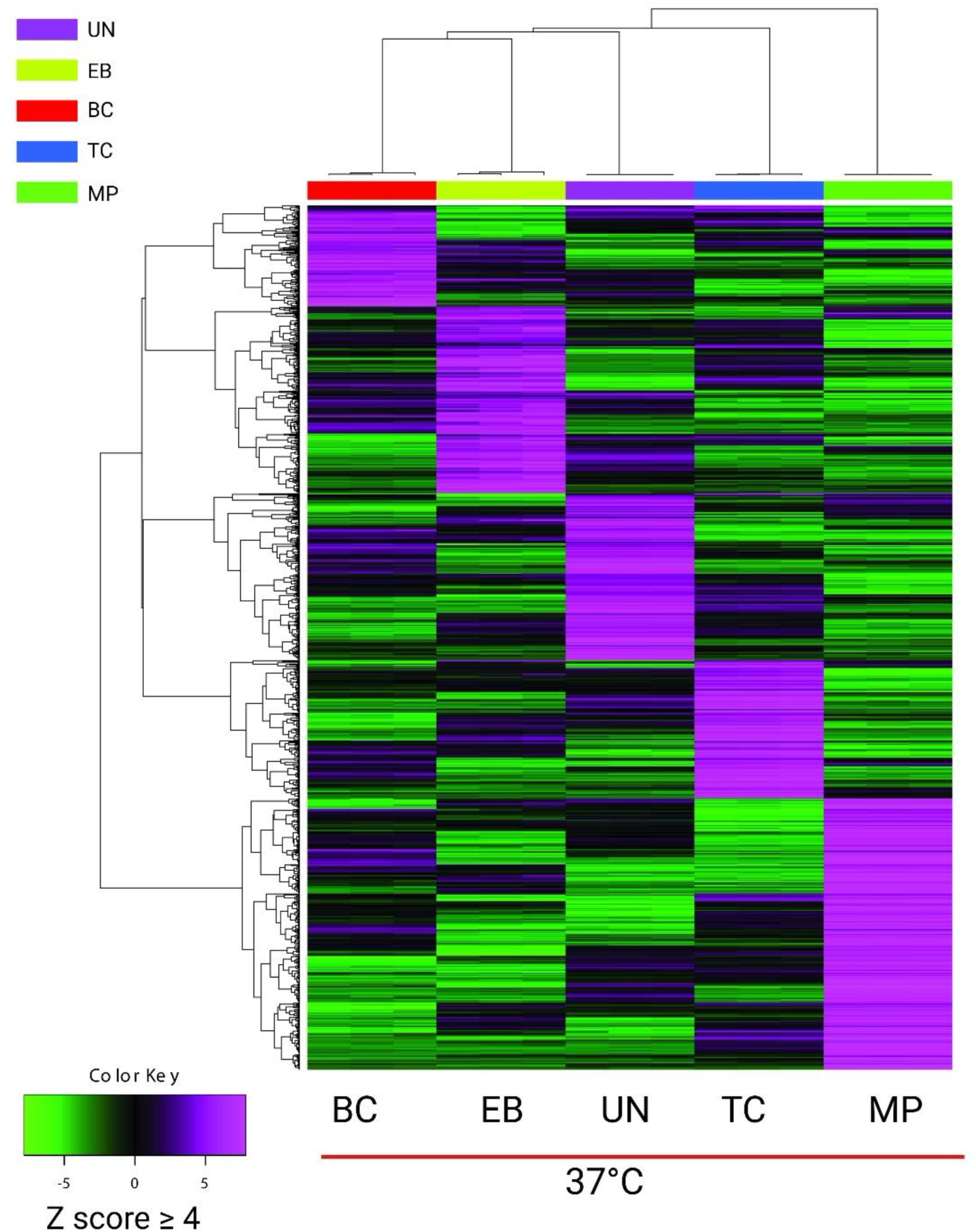
Hierarchical clustering heatmap showing gene expression across pollen developmental stages under heat stress: Uni-cellular (UN), early bi-cellular (EB), bi-cellular (BC), tri-cellular (TC), and mature pollen (MP). Sample groups are color-coded as indicated in the top left legend. Expression levels are represented by Z- scores, with a cutoff of ±4, where purple indicates up-regulation (Z ≥ 4) and green shows down-regulation (Z ≤ -4).

**Figure S2.**
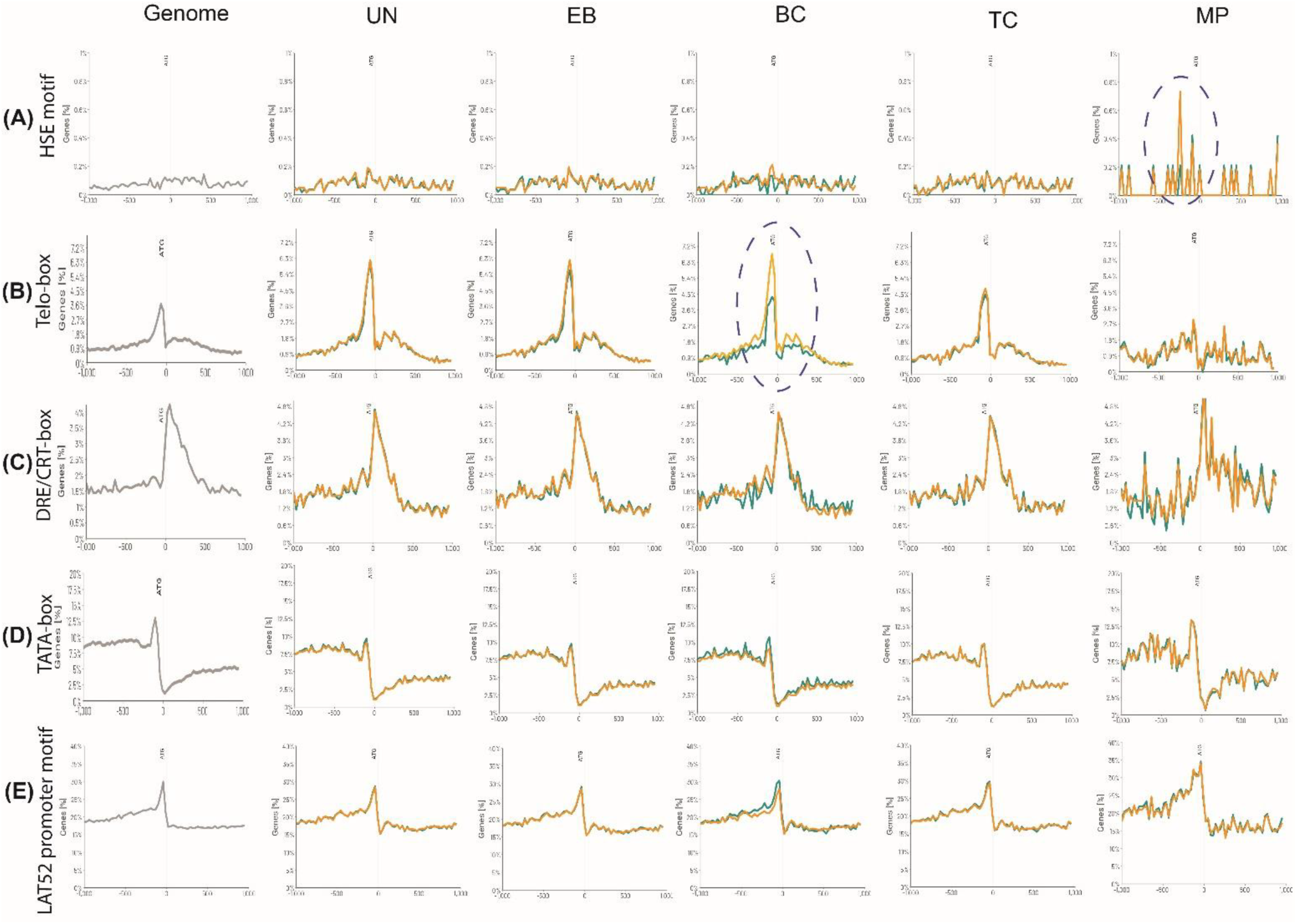
Localization and distribution of gene regulatory motifs in vicinity of ATG in genes highly expressed across pollen developmental stages under heat stress. (**a**) HSE motif (nGAAnnTTCnnGAAn), *Telo*-box (TAGGGTTT), (**c**) DRE/CRT-box (CCGAC), (**d**) TATA-box and (**e**) LAT52 promoter motif, were analyzed by the GOLEM program (https://golem.ncbr.muni.cz) in the promoters of the genes with the highest transcriptional activity during male gametophyte development. The developmental stages included: uni-cellular (UN), early bicellular (EB), bicellular (BC), tri-cellular (TC), mature pollen (MP) Motif distribution across the genome (grey) was compared to their occurrence in genes highly expressed under control condition (green) and heat stress (yellow). The genes expressed in the 90th percentile are shown within the range <-1000, 1000> bp. The axis size was adjusted in each row.

**Figure S3.**
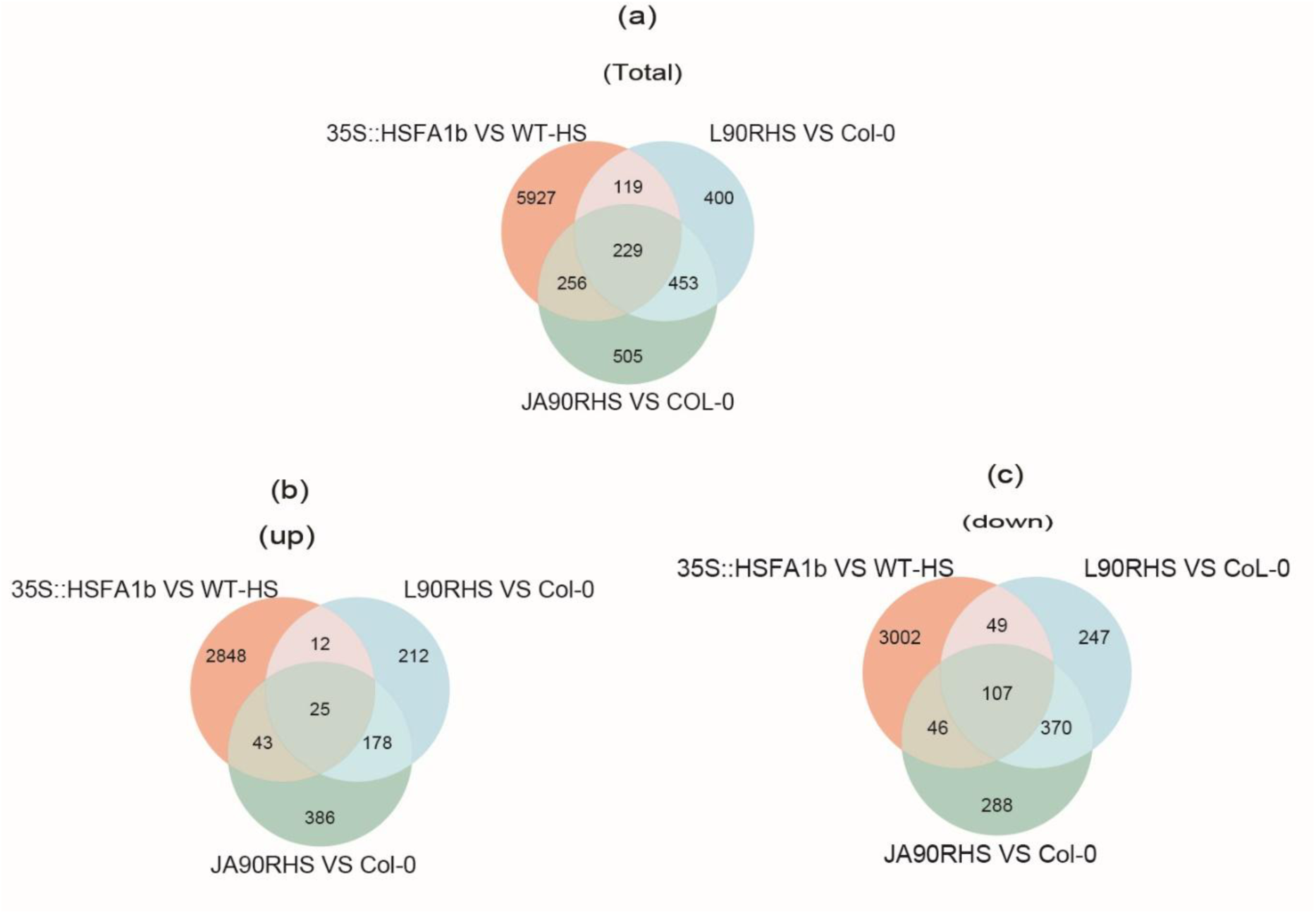
Comparative analysis of differentially expressed genes (DEGs) in heat stressed pollen of L90R and JA90R compared with DEGs in 35S:HSFA1b vs heat stressed WT in sporophytic tissue (leaves) (**a**) The Venn diagram shows the overlap in total DEGs among 35S::HSFA1 vs. WT under heat stress (red), L90R vs. WT (blue), and JA90R vs. WT (green). (**b**) The Venn diagram shows the overlap in up-regulated DEGs among 35S::HSFA1 vs. WT under heat stress (red), L90R vs. WT (blue), and JA90R vs. WT (green). (b) The Venn diagram shows the overlap in down-regulated DEGs among 35S::HSFA1 vs. WT under heat stress (red), L90R vs. WT (blue), and JA90R vs. WT (green).

**Figure S4.**
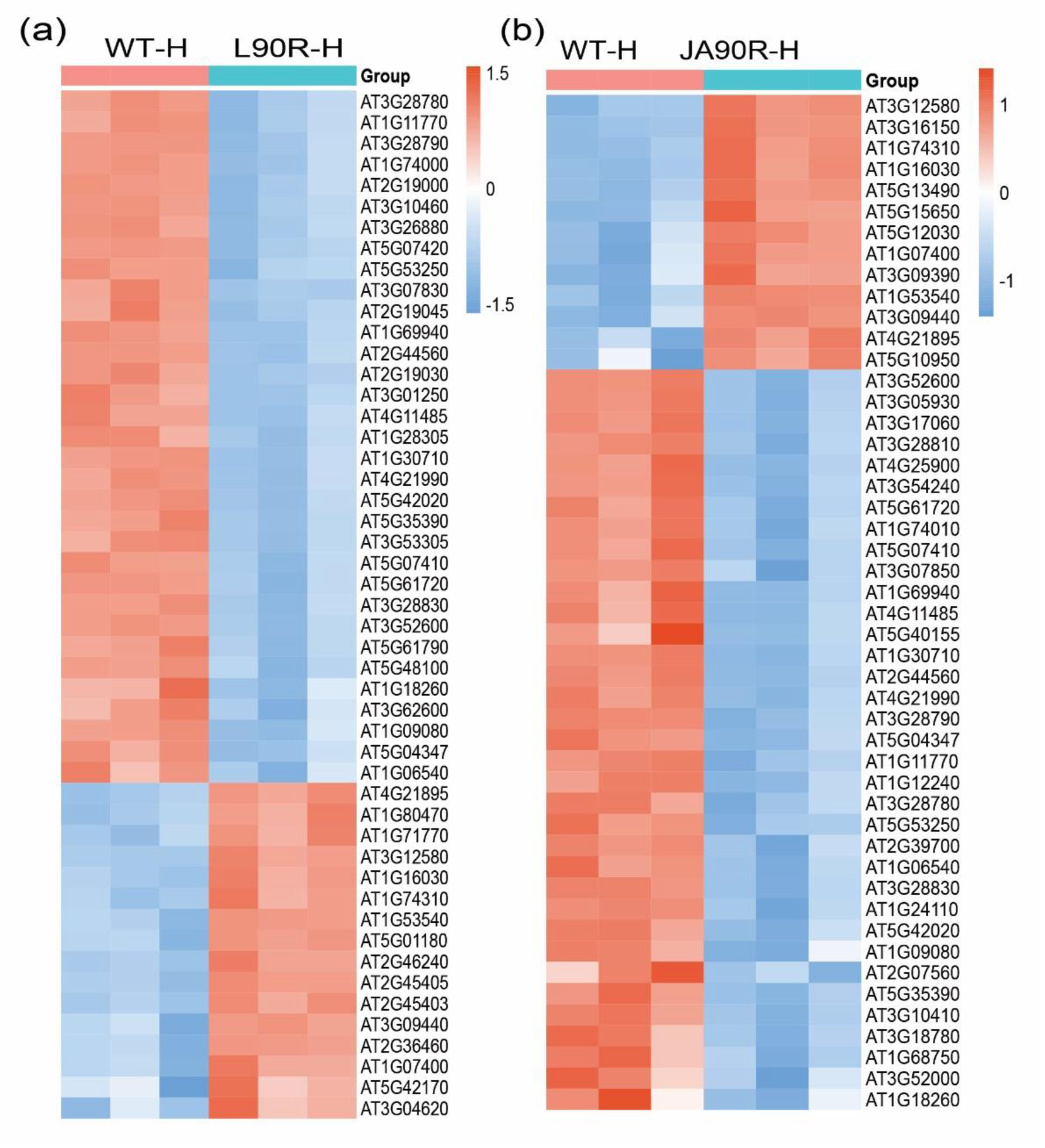
Comparative analysis of top 50 differentially expressed genes (DEGs) in heat stressed pollen of L90R and JA90R compared with heat stressed WT pollen. **(a)** Heatmap of top 50 DEGs comparing WT and L90R under heat stress (37°C), (**b**) Heatmap of top 50 DEGs comparing WT and L90R under heat stress (37°C), showing the expression levels of genes. Orange indicates up-regulation, whereas blue indicates down-regulation.

**Figure S5.**
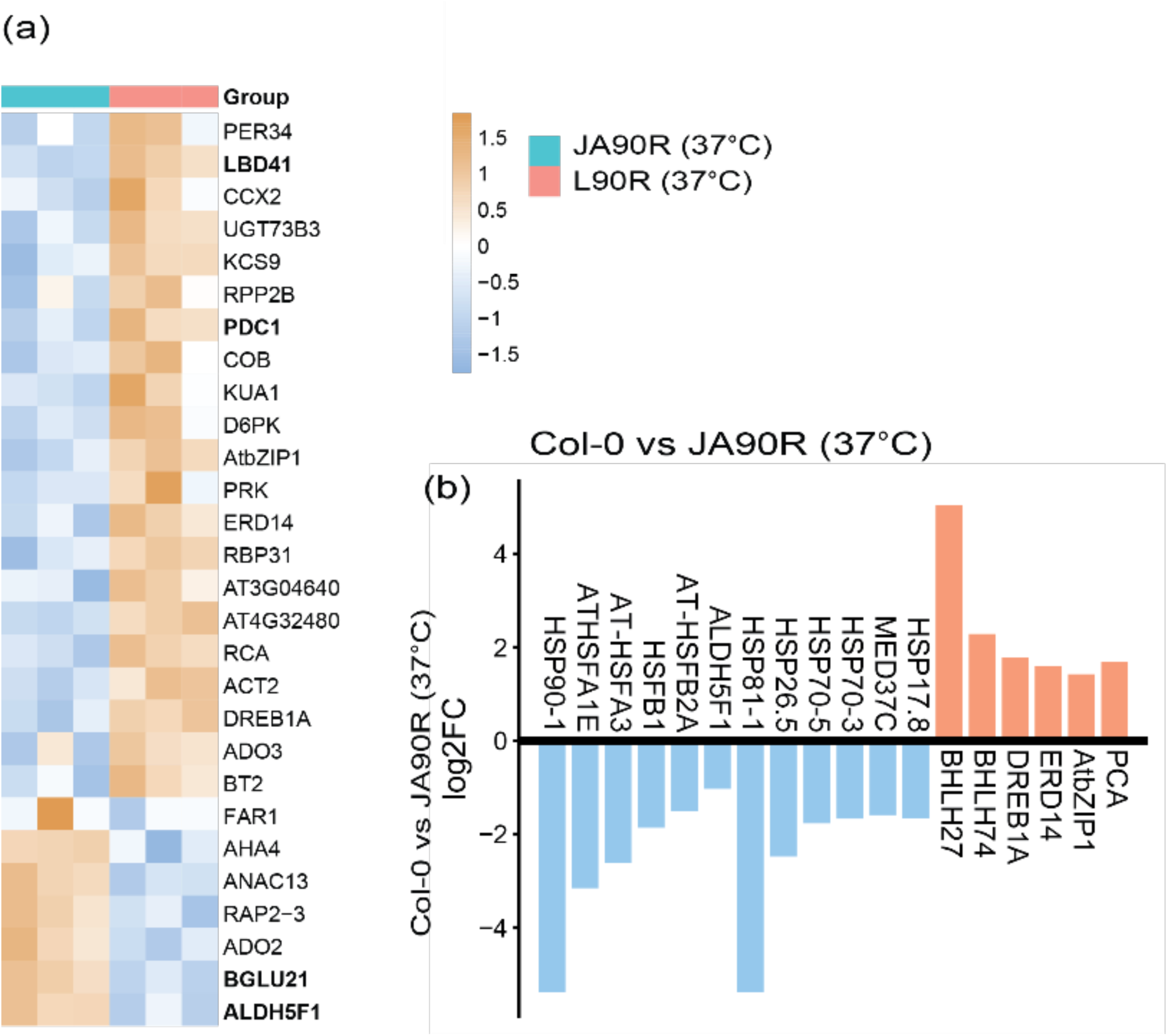
(a) Heatmap of heat stress-related genes, which are differentially expressed in JA90R vs L90R under heat stress conditions **(b)** log₂(Fold Change) values for selected genes involved in heat stress response in WT vs JA90R under heat stress (37°C). Blue bars represent down-regulated genes, while orange bars represen up-regulated genes.

**Table S1.**
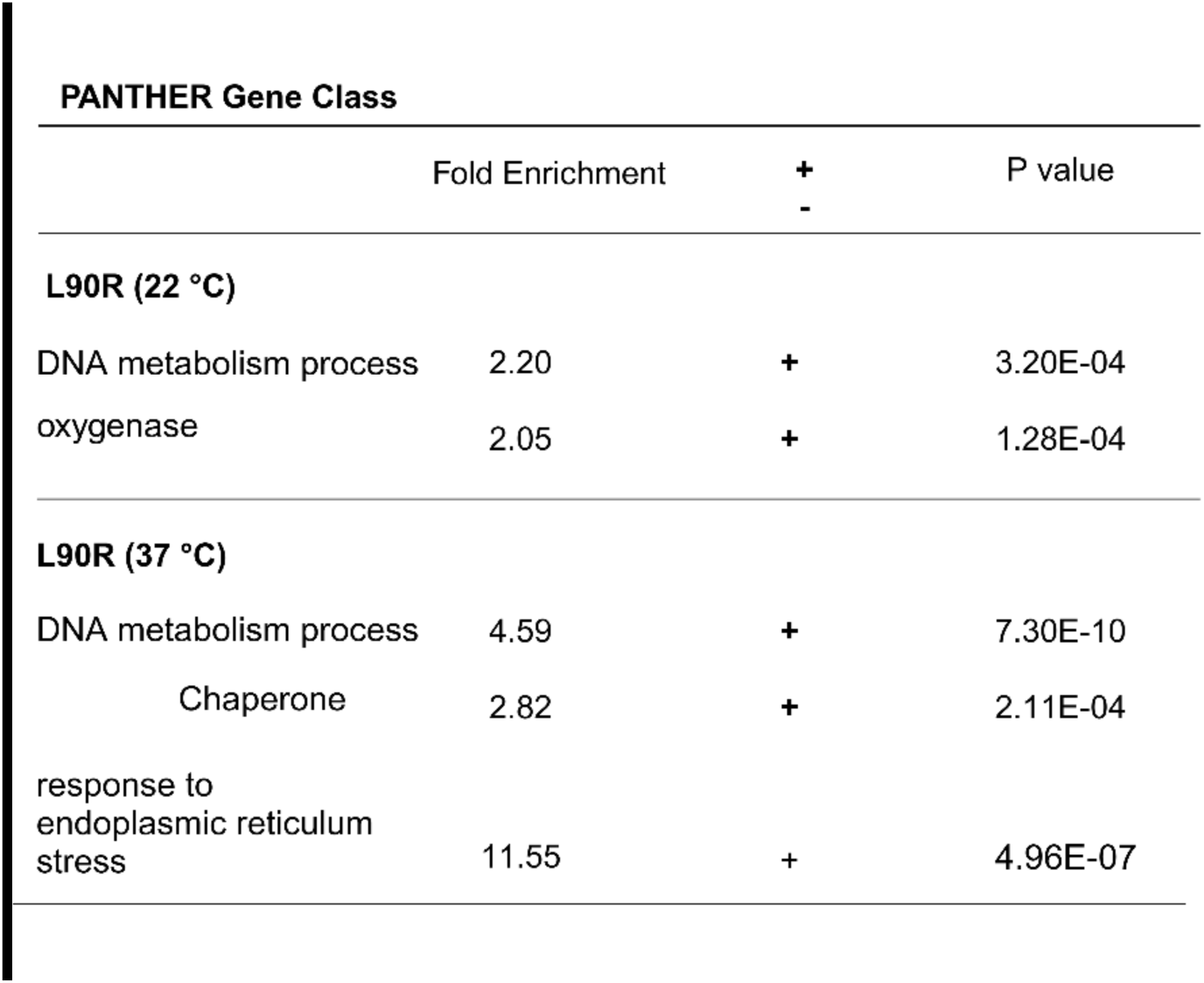
The down-regulation of genes involved in DNA metabolism and chaperone activity in mature pollen of L90R compared to WT is depicted. Differential gene expression analysis highlights significant reductions in the expression levels of these functional gene categories, suggesting a potential impact on DNA repair mechanisms and protein folding capacity in mature pollen in L90R lines

| PANTHER Gene Class |  |  |  |
| --- | --- | --- | --- |
|  | Fold Enrichment | +<br>- | P value |
| **L90R (22 °C)** |  |  |  |
| DNA metabolism process | 2.20 | + | 3.20E-04 |
| oxygenase | 2.05 | + | 1.28E-04 |
| **L90R (37 °C)** |  |  |  |
| DNA metabolism process | 4.59 | + | 7.30E-10 |
| Chaperone | 2.82 | + | 2.11E-04 |
| response to endoplasmic reticulum stress | 11.55 | + | 4.96E-07 |

**Supplementary data S1**. Gene onthology analysis for pollen stages under heat stress.

**Supplementary data S2.** DEGs in pollen stages under heat stress.

**Supplementary data S3.** DEGs in L90R and JA90R under heat stress.

**Supplementary data S4.** DEGs in 35S::HSFA1B under heat stress.

